# SAF-A promotes origin licensing and replication fork progression to ensure robust DNA replication

**DOI:** 10.1101/2021.03.22.436394

**Authors:** Caitlin Connolly, Saori Takahashi, Hisashi Miura, Ichiro Hiratani, Nick Gilbert, Anne D. Donaldson, Shin-ichiro Hiraga

## Abstract

The organisation of chromatin is closely intertwined with biological activities of chromosome domains, including transcription and DNA replication status. Scaffold attachment factor A (SAF-A), also known as Heteronuclear Ribonucleoprotein Protein U (HNRNPU), contributes to the formation of open chromatin structure. Here we demonstrate that SAF-A promotes the normal progression of DNA replication, and enables resumption of replication after inhibition. We report that cells depleted for SAF-A show reduced origin licensing in G1 phase, and consequently reduced origin activation frequency in S phase. Replication forks progress slowly in cells depleted for SAF-A, also contributing to reduced DNA synthesis rate. Single-cell replication timing analysis revealed that the boundaries between early- and late-replicating domains are blurred in cells depleted for SAF-A. Associated with these defects, SAF-A-depleted cells show elevated γH2A phosphorylation and tend to enter quiescence. Overall we find that SAF-A protein promotes robust DNA replication to ensure continuing cell proliferation.

## Introduction

DNA replication in eukaryotic genomes initiates from discrete sites termed DNA replication origins. Potential replication origin sites are defined by stepwise assembly of a protein complex, the pre-replication complex (pre-RC), during G1 phase of the cell cycle (Fragkos et al., 2015). During pre-RC formation the Origin Recognition Complex (ORC) and CDT1 cooperate to load the heterohexameric MCM complex, leading to ‘replication origin licensing’ (McIntosh and Blow, 2012). MCM plays a critical role when DNA replication initiates at each origin, forming the central component of the replicative helicase (Fragkos et al., 2015). Cells monitor the level of replication licensing to prevent cell cycle progression if an insufficient number of sites are licensed (Feng et al., 2003; Lau et al., 2009; Nevis et al., 2009; Shreeram et al., 2002; Zimmerman et al., 2013). This “licensing checkpoint” mechanism appears to be compromised or lost in cancer cells (Feng et al., 2003; Lau et al., 2009; Nevis et al., 2009; Shreeram et al., 2002; Zimmerman et al., 2013).

A recent study demonstrates that replication licensing is impacted by the state of chromatin packaging. The histone methyltransferase SET8 (also called PR-SET7 or KMT5A) can stimulate origin licensing at specific sites (Tardat et al., 2010), but also prevents over-licensing by enhancing chromatin compaction as cells exit mitosis (Shoaib et al., 2018). SET8 is responsible for the methylation of histone H4 at lysine 20, and for maintaining chromatin compaction at the M/G1 boundary (Shoaib et al., 2018). Replication licensing is therefore impacted both by local chromatin changes and broader changes occurring at chromosome domain level. Despite these discoveries, how chromatin packaging status affects origin licensing and the subsequent steps in DNA replication is still not fully understood.

There is however a well-established connection between chromatin packaging and the temporal programme of replication of chromosomal domains (Fu et al., 2018; Gilbert, 2010; Gilbert et al., 2010), in which euchromatin domains containing active genes generally replicate early in S phase, while heterochromatic, highly packaged domains containing mainly inactive genes replicate late. Replication timing of some domains is modulated during development, often reflecting changes in gene activity (Hiratani et al., 2008). The replication timing programme is established at early G1 phase, at a timing decision point (TDP) that coincides with chromatin decompaction and chromatin remodelling as cells exit M phase (Shoaib et al., 2018). The coincidence of this TDP with M/G1 phase suggests that the dynamic controls over chromatin structure imposed as cells exit mitosis determine the replication potential and subsequent replication timing of local chromatin domains (Dimitrova and Gilbert, 1999; Dimitrova et al., 2002).

Chromatin packaging also impacts on replication progression, with processive replication of heterochromatin regions requiring local decompaction (Chagin et al., 2019). Recent studies also highlight numerous “difficult-to-replicate” regions (Cortez, 2015; Gadaleta and Noguchi, 2017), including DNA-protein complexes, repetitive DNA such as centromeres and telomeres, and secondary DNA structures. Replicating such regions requires support by specific proteins, without which replication tends to fail, leading to genome instability (Cortez, 2015; Gadaleta and Noguchi, 2017) and the formation of fragile sites (Boteva et al., 2020). These observations highlight the importance of modulating chromatin structure during the replication process.

Scaffold Attachment Factor A (SAF-A; also known as Heterogeneous Nuclear Ribonucleoprotein U) is an RNA- and DNA-binding protein which modulates chromatin structure by tethering chromatin-associated RNA (caRNA) to chromatin (Fackelmayer et al., 1994; Kiledjian and Dreyfuss, 1992; Nozawa et al., 2017; Sharp et al., 2020). SAF-A oligomerisation contributes to de-compacted chromatin (Nozawa et al., 2017), and it has been shown that depletion of SAF-A causes global chromatin condensation (Fan et al., 2018). A super-resolution microscopy study also implicates SAF-A in the establishment of correct chromatin structure, and SAF-A has been shown to regulate both active chromatin and also X-chromosome inactivation (Smeets et al., 2014). SAF-A interacts and colocalises with proteins that define chromatin domain boundaries, namely CTCF and cohesin, and plays a role in defining boundaries of Topologically-Associated Domains that form smaller units of chromosome organisation (Fan et al., 2018; Zhang et al., 2019).

Association of SAF-A with chromatin is cell cycle-regulated: SAF-A is chromatin-associated throughout interphase but is removed from chromatin in M phase (Sharp et al., 2020). Regulated dissociation of SAF-A from the mitotic chromosome, triggered by phosphorylation of SAF-A by Aurora B protein kinase, is essential for proper progression of mitosis (Douglas et al., 2015; Sharp et al., 2020). SAF-A re-associates with chromatin as cells exit from mitosis, implicating SAF-A in chromatin decompaction at this cell cycle stage.

SAF-A also localises to DNA damage sites swiftly after γ-ray irradiation, and then at a later stage appears to be excluded from damage sites (Hegde et al., 2016). The molecular function of SAF-A in damage repair is yet to be demonstrated, but one possibility is that SAF-A quickly modifies local chromatin structure at damage sites to facilitate action of the repair machinery.

Interestingly, the expression of the SAF-A gene tends to increase in a wide range of cancers, particularly in breast invasive carcinoma (The Cancer Genome Atlas). This increased expression suggests that SAF-A contributes to the formation or survival of cancer cells in a dose-dependent manner. Conversely, SAF-A loss-of-function alleles are linked to developmental disorders including microcephaly (Durkin et al., 2020; Leduc et al., 2017; Yates et al., 2017). Overall, these observations suggest a positive role for SAF-A in promoting cell proliferation. While roles of SAF-A in mitosis have been investigated (Sharp et al., 2020), its contribution to cell proliferation during interphase, particularly to DNA replication, has not been studied.

Here we investigate the effects of SAF-A on DNA replication. We show that SAF-A protein is required for full replication licensing in the G1 phase of the cell cycle, and depleting SAF-A leads to increased spacing between replication origins. We find moreover that replication fork progression is impeded in cells depleted for SAF-A, and that SAF-A protein plays a role in defining the boundaries of early/late replication domains in the genome-wide replication programme. Loss of these functions leads to spontaneous replication stress and increases cellular entry to quiescence, explaining the need for SAF-A for normal cell proliferation.

## Results

### SAF-A is required for robust DNA replication

To assess the general impact of SAF-A on subnuclear organisation of chromatin, we examined the distribution of DNA within nuclei of hTERT-RPE1 cells treated with siRNA targeting SAF-A (siSAF-A) (Fig 1A). hTERT-RPE1 is a non-cancer cell line derived from retinal pigment epithelial cell immortalised by expression of human telomerase (hTERT) (Bodnar et al., 1998). Using super-resolution microscopy to examine very thin sections, we found that control nuclei show relatively homogeneous DNA density distribution (i.e. mostly green), with smaller areas of higher (yellow/red) or lower (cyan/blue) DNA density. In siSAF-A cells the DNA density distribution shows larger areas with low DNA density (blue) interspersed with high DNA density areas (red), indicative of a more polarised distribution with sections of genomic DNA densely packed in abnormally compact domains. Unbiased classification of “DAPI-positive” and “DAPI-negative” areas in each nucleus (Fig 1B & S1A) confirmed the formation of larger “DAPI-negative” areas in siSAF-A nuclei, with chromatin confined into smaller areas. These data suggest that SAF-A promotes properly dispersed chromatin distribution within nuclei, and plays a role in preventing the formation of over-compacted chromatin. This microscopic observation is consistent with SAF-A function in maintaining correct chromatin architecture as revealed by microscopy (Nozawa et al., 2017) and Hi-C methods (Fan et al., 2018).

**Figure 1:**
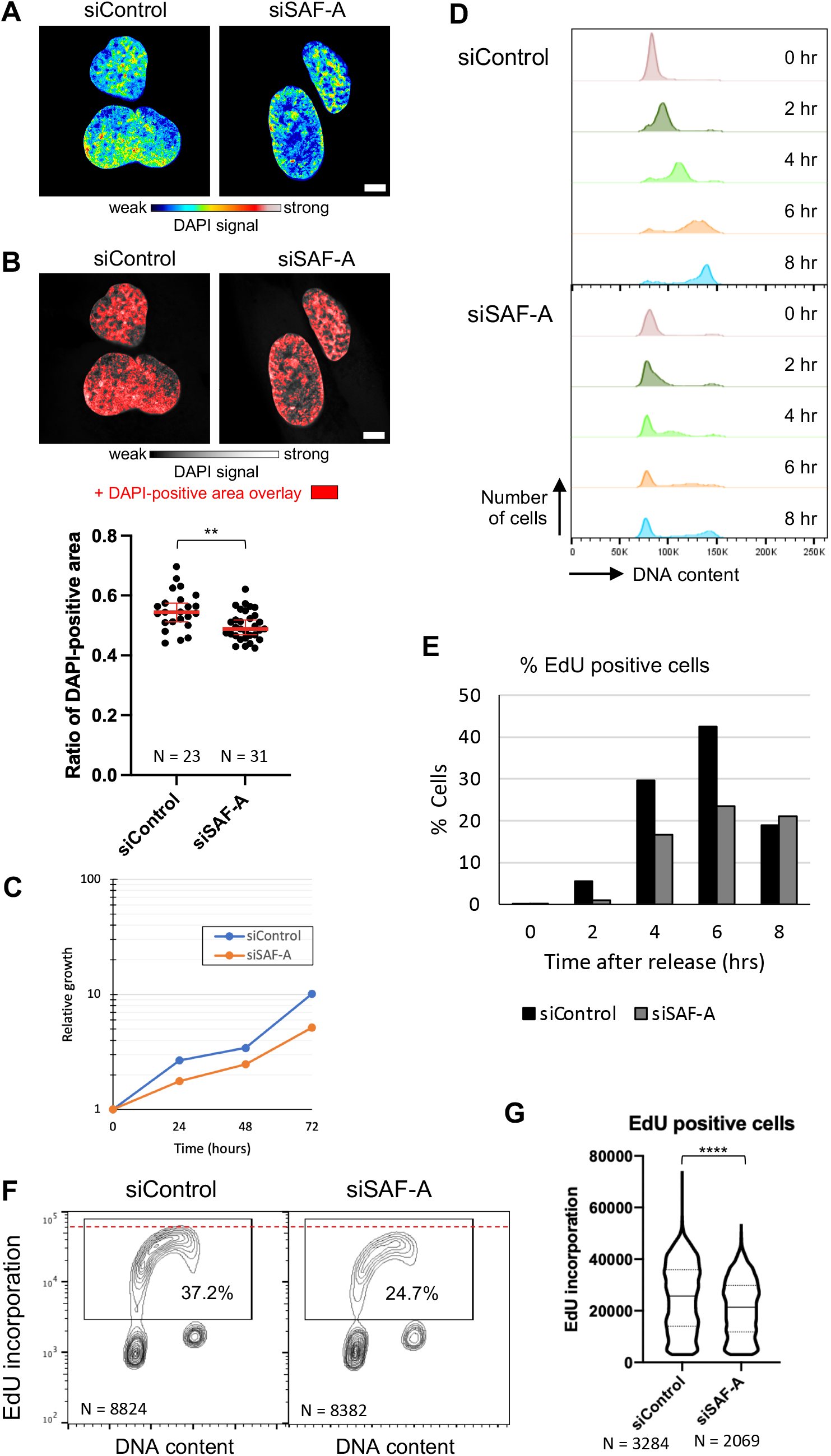
SAF-A is required for robust DNA replication. (A) Specimen images showing the distribution of DNA within nuclei of cells treated with control siRNA (siControl) and SAF-A siRNA (siSAF-A). Super-resolution images of DAPI-stained DNA were identically processed and are shown in a pseudo-colour scale as shown below images. Bar is 5 μm. (B) Quantification of “DAPI-positive” areas. Images were analysed by an automated pipeline and the ratio of “DAPI-positive” area relative to the entire nucleus calculated at the middle section of each nucleus. Images at top show DAPI intensity in greyscale with identified “DAPI-positive” areas overlaid in red. Plots below show the distribution of the “DAPI-positive” proportions of nuclei; median with 95% confidence interval shown in red. Statistical significance was calculated by Student’s t-test. N=23 for siControl and 31 for siSAF-A. (C) Growth of hTERT-RPE1 cells in DMEM-F12 media treated with control siRNA (siControl) or SAF-A siRNA (siSAF-A) was measured by counting the number of cells at each passage. (D) Cells depleted for SAF-A are defective in recovery from replication stress. siControl and siSAF-A cells were arrested with 4 mM HU for 24 hr, and released into fresh media. Cells were pulse-labelled with 20 μM EdU for 15 min before sampling at indicated time points. Cells were fixed, differential labelled with CellTrace Yellow, and analysed for DNA content and EdU incorporation by flow cytometry. Contour lines are set to 5% so that 5% of cells fall between each pair of contour lines. (E) Percentages of EdU-positive cells after removal of HU. After removal of HU, cells were pulse-labelled for 20 min with EdU at the indicated time points and EdU-positive cells were identified by flow cytometry. See Fig S1C for gating strategy. (F) Cells depleted for SAF-A show reduced DNA synthesis rate. Asynchronously growing cells were pulse-labelled with 20 μM EdU for 1 hr and collected. DNA content and the amount of EdU was measured by flow cytometry. (G) Incorporation of EdU in S phase cells. EdU incorporation per cell was measured in EdU-positive (S phase) cells. Violin plots show the median (solid line) and quartiles (dotted lines). As the distribution was not normal, the statistical significance was calculated using Mann-Whitney-Wilcoxon test. ** p ≤ 0.01, **** p ≤ 0.0001.

Cells depleted for SAF-A were reported to show proliferation defects (Nozawa et al., 2017), but the exact nature of the defect was not studied in detail. We examined the cell proliferation and DNA replication profiles of cells depleted for SAF-A. siSAF-A cells showed a significant and reproducible reduction in cell proliferation rates, compared with siControl cells (Fig 1C), consistent with a previous report (Nozawa et al., 2017). Flow cytometry analysis of DNA content in asynchronous cultures (Fig S1B) however showed no specific cell cycle arrest point, but did reveal a slight reduction of S phase population, suggesting that loss of SAF-A may cause problems with DNA replication.

Cells depleted for SAF-A were reported to be defective in recovery from replication inhibition by the DNA polymerase inhibitor aphidicolin (Nozawa et al., 2017). We therefore tested whether SAF-A-depleted cells also fail to recover from the DNA replication inhibitor hydroxyurea (HU), an inhibitor of ribonucleotide reductases that causes stalled replication forks. After treating siControl and siSAF-A cells with 4 mM HU for 24 hr to cause early S phase arrest (Fig 1D, 0 hr), we examined recovery by monitoring DNA content and incorporation of a thymidine analogue ethynyl deoxyuridine (EdU). Control cells recovered from arrest efficiently and reached mid-S phase by 4 hr after release from HU (Fig 1D, siControl). In contrast, very few siSAF-A cells recovered to reach a similar stage by 6 hr (Fig 1D, siSAF-A). Assessment of EdU-positive cells further indicated that a reduced number of siSAF-A cells were able to resume DNA synthesis compared with siControl (Fig 1E), and that the rate of DNA synthesised in EdU-positive siSAF-A cells was lower than that in siControl cells until 6 hr after release (Fig S1C & D). By 8 hr after the release, the majority of siControl cells had finished DNA replication, whereas a notable fraction of siSAF-A cells were still synthesising DNA (Fig S1C & D). These observations indicate that cells depleted for SAF-A have difficulty in recovering from HU. Taken together, deficiency of SAF-A causes cells to be severely impaired in recovery from replication stress.

We tested whether depletion of SAF-A impacts DNA replication in the absence of exogenous stress, by measuring DNA synthesis rate based on pulse-labelling nascent DNA with EdU followed by flow cytometry analysis. Cells depleted for SAF-A showed a significantly reduced percentage of EdU-positive cells (Fig 1F; 37.2% in siControl and 24.7% in siSAF-A). Moreover, the EdU-positive population of siSAF-A cells showed reduced DNA synthesis rate compared to siControl (Fig 1G). This DNA synthesis defect of siSAF-A cells is not confined to a specific stage of S phase (Fig S1E), suggesting that SAF-A function is required throughout DNA replication.

Together, these results indicate that SAF-A is required for robust DNA replication without exogenous replication stress, and also supports the recovery of cells after replication stress.

### SAF-A is important for replication licensing

Changes in chromatin due to SAF-A depletion could potentially affect multiple steps of DNA replication, including origin licensing, replication fork progression, and fork restart. We decided to assess the requirement for SAF-A for each of these steps in a series of experiments.

Since SAF-A plays a positive role in open chromatin structure, and prevents over-compaction (Fig 1A, 1B, and S1A) we hypothesised that SAF-A may play a positive role in stimulating origin licensing by promoting open chromatin. To test this hypothesis, we used a flow cytometry ‘3D licensing assay’ (Moreno et al., 2016) (Fig 2A & B), which allows simultaneous measurement of MCM loading on chromatin and incorporated EdU (to assess cell cycle stage). In this assay (Fig 2A) G1 (red box), S (cyan), and G2/M (orange) phase cells can be clearly distinguished, and the amount of chromatin-associated MCM3 assessed in each cell cycle population (Fig 2B). As clearly seen in Fig 2B, siSAF-A cells show reduced levels of chromatin-associated MCM3 in individual cells both in G1 phase (red) and in cells entering S phase (left hand part of cyan population). This observation indicates that siSAF-A cells show compromised levels of MCM loading in G1 phase cells that persist into S phase, suggestive of a defect in origin licensing.

**Figure 2:**
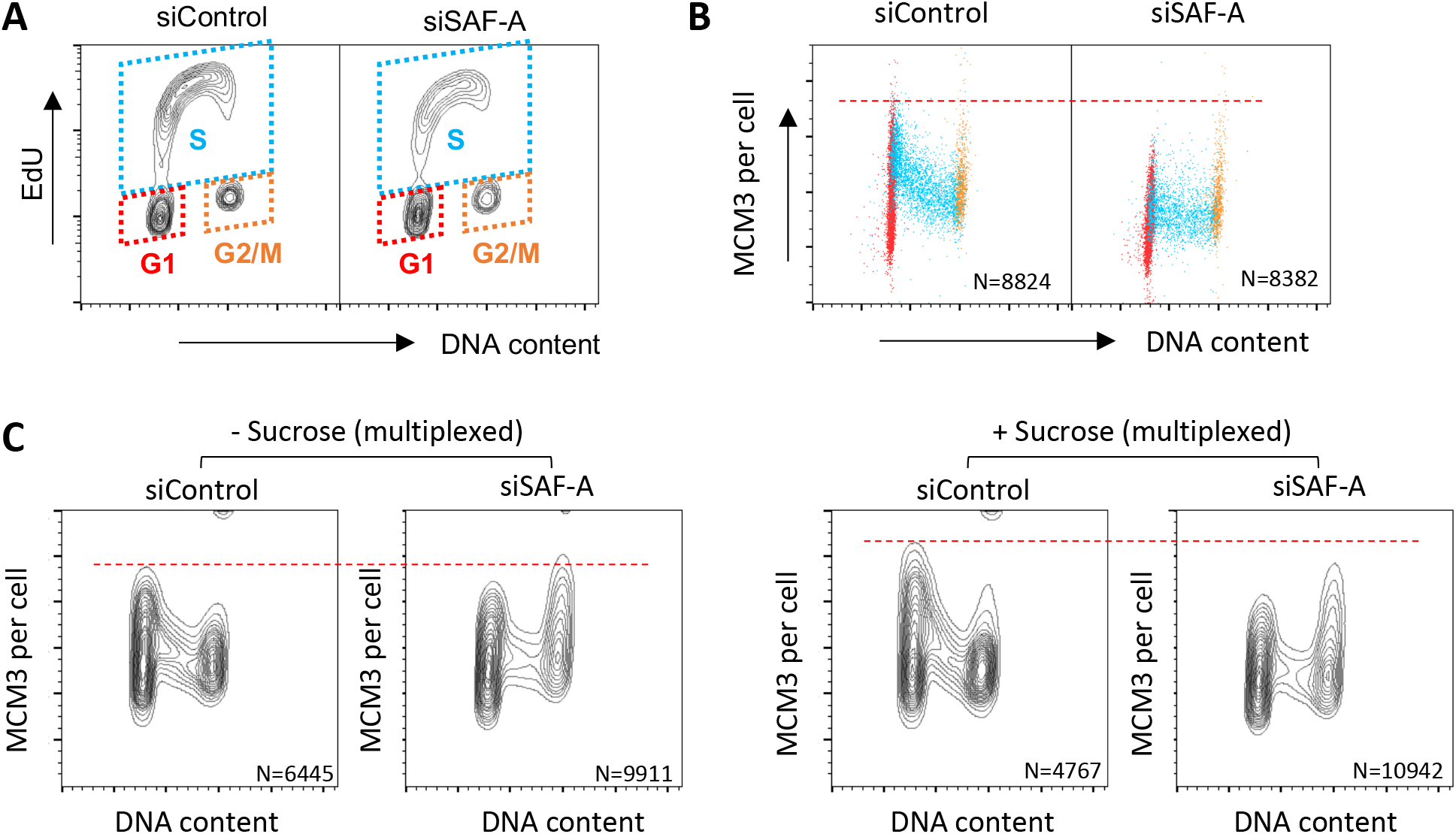
SAF-A is important for replication licensing. (A) Separation of cell cycle phases of hTERT-RPE1 cells by DNA content and EdU incorporation. Gates used in (B) are indicated by coloured dotted parallelograms. Contour lines are set to 5% so that 5% of cells fall between each pair of contour lines. (B) 3-D licensing assay in hTERT-RPE1 cells. G1 (red), S (Cyan), and G2/M (Orange) cell populations were distinguished as in (A). Chromatin-associated MCM3 was measured as previously described (Hiraga et al., 2017). (C) SAF-A is required to protect licensing against chromatin over-compaction. The effect of depleting SAF-A on licensing in hTERT-RPE1 cells was tested without or with Sucrose, which induces chromatin over-compaction. Cells were treated with 80 mM sucrose for 5 hrs before fixation. MCM3 chromatin association was tested as in (B), but without EdU and with “multiplexing” (see Materials and Methods). Note that siControl and siSAF-A samples are 2-way multiplexed (Both −Sucrose samples together or both +Sucrose samples together), so that siControl and siSAF-A samples are quantitatively comparable, but −Sucrose and +Sucrose samples are not. Contour lines are set to 5%.

Treatment of cells with sucrose is known to induce chromatin over-compaction, through a molecular crowding effect (Richter et al., 2007). This sucrose-induced over-compaction has been shown to compromise origin licensing (Shoaib et al., 2018). We found that the combination of sucrose treatment and SAF-depletion (Fig 2C right) enhanced the licensing defect caused by SAF-A depletion (Fig 2C left). This result indicates that SAF-A counteracts chromatin compaction to promote replication licensing. Note that increased chromatin association of MCM3 in G2/M cells is occasionally observed but not consistently caused by SAF-A depletion.

Taken together, these data demonstrate that SAF-A is required for normal levels of MCM loading, and helps the licensing machinery to resist chromatin over-compaction.

### SAF-A affects other replication licensing factors

We next tested whether SAF-A affects other licensing proteins. CDT1 interacts with the MCM complex and assists its loading onto chromatin in G1 phase (Frigola et al., 2017; Zhai et al., 2017a): outside of G1 phase CDT1 is negatively regulated by Geminin (Blow and Tanaka, 2005). *In vitro* studies show that CDT1 dissociates from MCM after the assembly of the double MCM hexamer on DNA (Zhai et al., 2017b). Consistently, flow cytometry analysis showed that CDT1 associates with chromatin predominantly in G1 phase (vertical spikes in Fig 3A). We found that depletion of SAF-A in hTERT-RPE1 cells caused reduced chromatin association of CDT1 (Fig 3A, siSAF-A), consistent with the reduced MCM licensing in G1 phase (Fig 2B).

**Figure 3:**
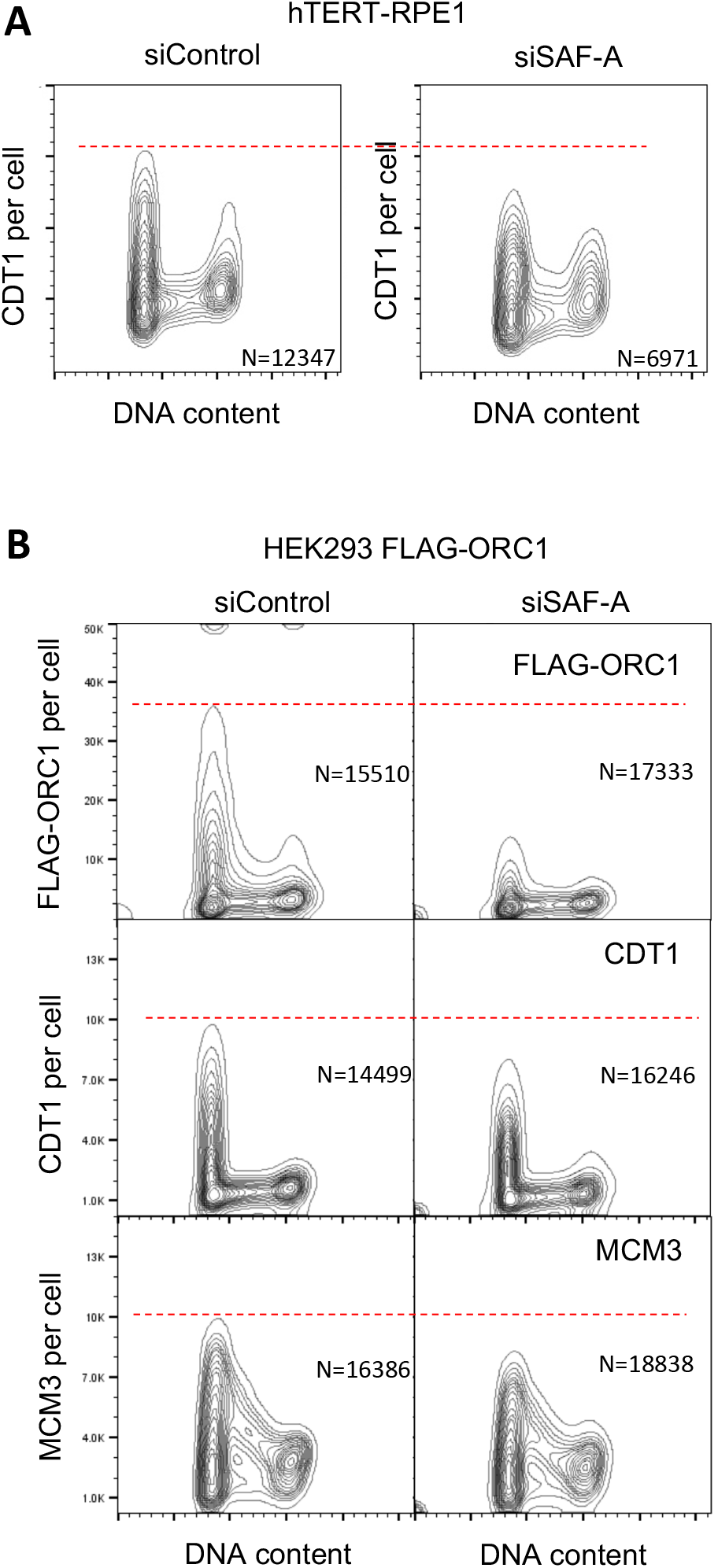
SAF-A affects multiple replication licensing factors. (A) SAF-A promotes chromatin association of CDT1 protein in G1 phase. Chromatin association of CDT1 protein in control and SAF-A depleted cells was tested in hTERT-RPE1 cells as in Fig 2C. Contour lines are set to 5% so that 5% of cells fall between each pair of contour lines. (B) SAF-A is required for full chromatin association of ORC1 and CDT1 proteins in G1 phase. Chromatin association of FLAG-tagged ORC1 protein in a HEK293-derived cell line was tested using anti-FLAG antibody (top panels). Chromatin association of CDT1 (middle panels) and MCM3 (bottom panels) proteins was tested in the same batch of cells. Contour lines are set to 5%.

We next tested the chromatin association of ORC1 protein. ORC1 protein is a subunit of the ORC protein complex that initially defines MCM loading sites. ORC1 protein expression and stability is cell cycle-regulated so that it is present predominantly in G1 phase, helping to confine MCM loading to G1 phase of the cell cycle (Mendez et al., 2002; Ohta et al., 2003; Tatsumi et al., 2003). In the absence of an ORC1 antibody suitable for flow cytometry analysis, we made use of a HEK293-based cell line expressing FLAG-tagged ORC1 protein (Tatsumi et al., 2003) to analyse chromatin association of ORC1 (Fig 3B). As expected and previously reported (Hiraga et al., 2017), ORC1 chromatin association is detected predominantly in the G1 phase of the cell cycle (Fig 3B, top panel). Depletion of SAF-A results in a reduction of chromatin-associated ORC1 (Fig 3B, top panel, right). We also observed that CDT1 and MCM licensing are reduced when SAF-A is depleted in the HEK293 FLAG-ORC1 cell line (Fig 3B, middle and bottom panels, respectively), similar to the effects in hTERT-RPE1 cells (Fig 2B & 3A). Loading of CDT1 was also hypersensitive to the combination of sucrose treatment and SAF-A depletion in HEK293 FLAG-ORC1 cells (Fig S2A). In contrast, chromatin association of ORC2, which does not fluctuate during cell cycle (Mendez and Stillman, 2000; Mendez et al., 2002), was not affected by SAF-A depletion (Fig S2B).

Reduced association of licensing factors in SAF-A-depleted hTERT-RPE1 cells was confirmed by Western analysis of chromatin-associated proteins. Western blotting of chromatin-enriched fractions confirmed the reduced association of CDT1 protein (Fig S2C, lanes 3 & 4) and ORC1 protein (Fig S2C, lanes 7 & 8 top panel) after SAF-A depletion. CDC6, another protein required for replication licensing (Blow and Tanaka, 2005), however did not show such a reduction in chromatin association (Fig S2C, lanes 7 & 8 middle panel). We did not assess CDC6 chromatin association by flow cytometry, because commercially available antibodies tested were unsuitable for flow cytometry (data not shown).

Overall, the data presented show that SAF-A promotes the G1-specific chromatin association of several origin licensing components, including loading of the MCM complex itself.

### SAF-A depletion results in reduced origin activation

Impaired replication licensing in cells depleted for SAF-A suggests there will be a reduced number of potential replication origins available for activation. Therefore, we next tested whether a reduced origin frequency is observed on chromosomes in SAF-A depleted cells, by measuring inter-origin distances using single-molecule DNA fibre analysis.

To detect origin activation on single DNA molecules, nascent DNA was sequentially labelled with thymidine analogues 5-chloro-2’-deoxyuridine (CldU) and 5-Iodo-2’-deoxyuridine (IdU), as illustrated in Fig 4A. Analogue incorporation was analysed by immunostaining of DNA fibres stretched by molecular DNA combing (Bianco et al., 2012). Replication origins can be identified as illustrated in the top panel of Fig 4B, with the mid-point between divergent replication forks assigned as a replication origin. In these experiments, increased distance between replication origins (inter-origin distances; IOD) is indicative of fewer active origins. We found that depletion of SAF-A caused an increase in IOD compared with the control (Fig 4B), suggesting that the number of active origins is indeed reduced by SAF-A depletion. This reduction in origin activation frequency probably reflects inefficient origin licensing.

**Figure 4:**
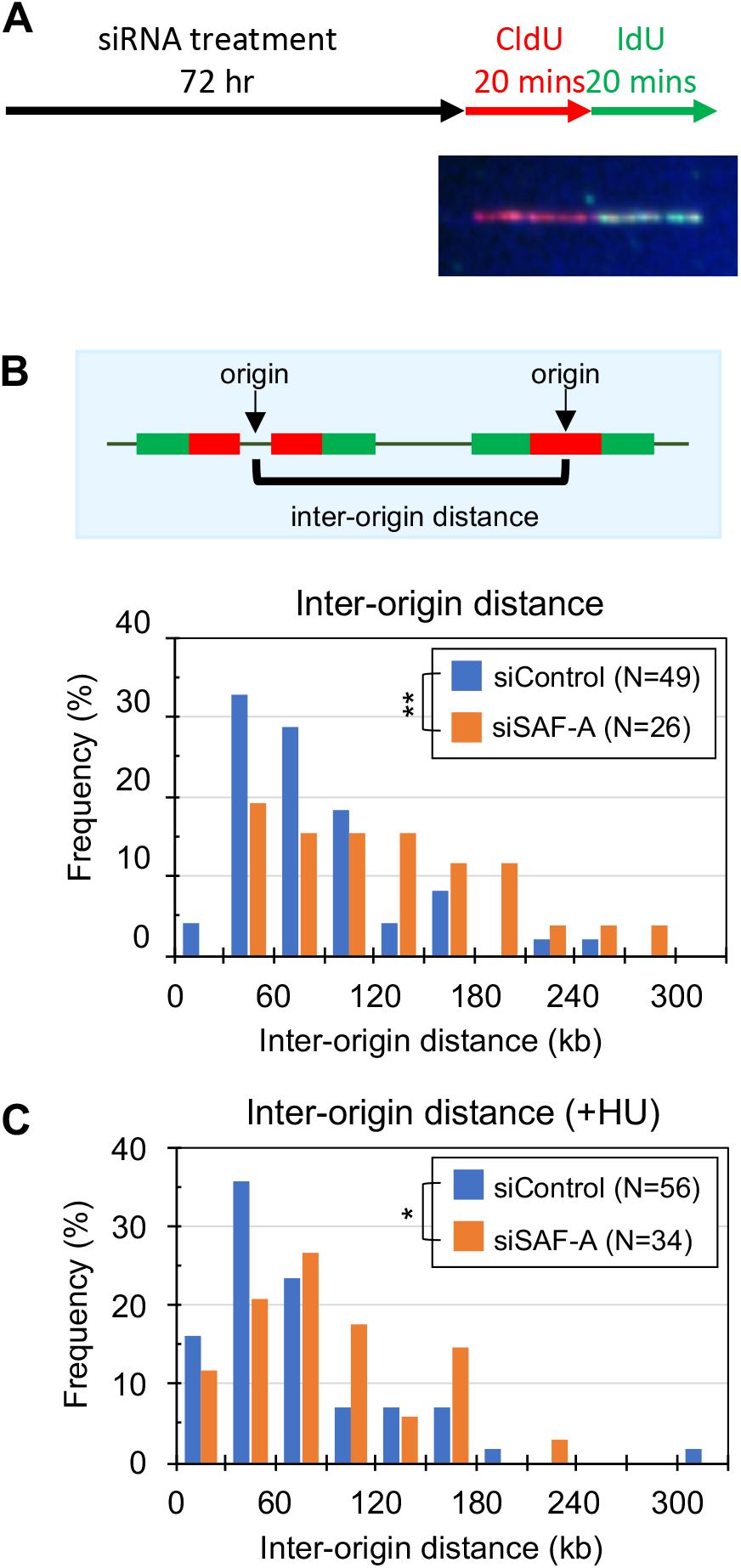
Cells depleted for SAF-A has a reduced origin activation potential and is defective in the activation of dormant origins. (A) Scheme of experiment. Cells were treated with either control siRNA (siControl) or SAF-A siRNA (siSAF-A) for 72 hr and sequentially pulse-labelled with CldU and IdU for 20 min, respectively. Cells were collected, and genomic DNA subjected to DNA combing. Specimen image shows visualised CldU and IdU. (B) Inter-origin distance was measured as illustrated in cells treated with control siRNA or SAF-A siRNA. The difference is statistically significant (p-value = 0.0066; Mann-Whitney-Wilcoxon test). (C) Inter-origin distance was measured in HU-treated cells. CldU and IdU labeling was carried out at the end of 4 hr HU treatment. The difference is statistically significant (p-value = 0.0353; Mann-Whitney-Wilcoxon test). Note that the HU concentration in this experiment is 0.1 mM, which does not stop DNA synthesis completely (Fig S3).

Stalling of DNA replication forks due to replication stress causes activation of nearby dormant origins (Ge et al., 2007), believed to protect cells from replication stress by guarding against the formation of unreplicated stretches between two stalled or collapsed replication forks (Blow and Ge, 2009; Kawabata et al., 2011). Since the licensing defect of SAF-A-depleted cells might affect the number of available dormant origins, we assessed whether cells depleted for SAF-A activate dormant origins normally (Fig 4C). siRNA-treated cells were incubated for 4 hr with 0.1 mM HU to slow replication forks. At the end of the HU treatment, nascent DNA was labelled with CldU and IdU as in Fig 4A. Under this condition, the replication fork speed is significantly reduced but still detectable by DNA combing (Fig S3). As expected, HU treatment induced the activation of dormant origins near stalled forks, evidenced by a leftward shift of overall IOD distribution and the appearance of very short IODs below 30 kb, both in siControl and siSAF-A cells (Fig 4C). In cells depleted for SAF-A, however, short IODs (in the range 0-30 kb) were observed at reduced frequency compared to siControl, suggesting an impaired dormant origin activation.

Overall these data confirm that the number of active DNA replication origins is reduced in cells depleted for SAF-A, and that SAF-A is required for activation of dormant origins at normal frequency under replication stress.

### SAF-A supports DNA replication fork progression

SAF-A depletion leads to decreased cellular DNA synthesis rate in unperturbed S phase (Fig 1F & G), as well as reduced origin licensing (Fig 2 & 3) and activation (Fig 4). However, it was unclear whether reduced origin activation fully accounts for the decreased cellular DNA synthesis. To explore whether altered DNA replication fork speed also affects DNA synthesis when SAF-A is deficient, we investigated replication fork speed using the same DNA combing technique. The lengths of IdU tracts in stretched DNA molecules were taken as a proxy for DNA synthesis rate. We find that replication fork speed is somewhat reduced in cells depleted for SAF-A (Fig 5A siSAF-A) compared with control cells (Fig 5A, siControl), indicating that SAF-A function is required for the normal rate of DNA replication fork progression.

**Figure 5:**
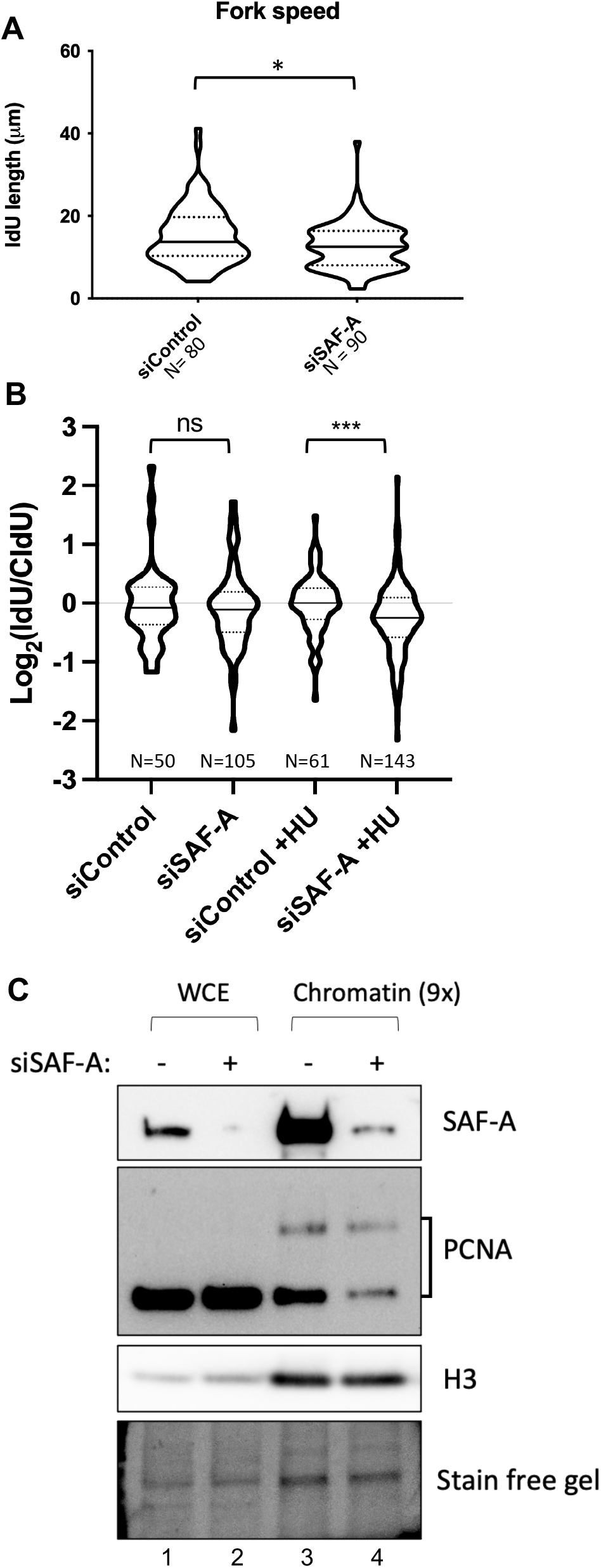
SAF-A supports replication fork progression and origin activation. (A) SAF-A is required to support normal replication fork speed. Nascent DNA was labelled as in Fig 4A, and replication fork speed measured based on the IdU tract length. Violin plots show the median (solid line) and quartiles (dotted lines). As distribution was not normal, statistical significance was tested by Mann-Whitney-Wilcoxon test. (B) SAF-A is required for fork processivity. The ratio of IdU:CldU tract length was measured in each where they appeared consecutive, and log base 2 values are plotted. Violin plots show the median (solid line) and quartiles (dotted lines). Statistical significance was tested by Student’s t-test (two-tails, unequal variance). (C) Chromatin association of molecular clamp PCNA is significantly reduced in cells depleted for SAF-A. Chromatin association of PCNA protein was assessed by chromatin fractionation. The abundance of SAF-A and PCNA in the chromatin-enriched fraction was assessed by western blotting. Histone H3 blot and Stain Free gel images shown as loading control. The upper PCNA band observed in the chromatin fractions probably corresponds to a SUMOylated form (Gali et al., 2012). ns, not significant; *p ≤ 0.05, ***p ≤ 0.001.

We next tested whether SAF-A depletion affects the processivity of DNA replication forks. When the processivity of DNA replication is high, the rate of DNA synthesis should be consistent through the CldU and IdU labelling periods, and the log value of (IdU tract length / CldU tract length) is expected to be close to 0. Frequent pause or collapse of forks would lead to a wider spread in log_2_(IdU /CldU) values. In unperturbed conditions, both siControl and siSAF-A cells indeed show median log_2_(IdU / CldU) values close to 0 (Fig 5B, left half). In contrast, when replication forks were challenged by HU, siSAF-A cells showed decreased log_2_(IdU / CldU) values (Fig 5B, compare ‘siSAF-A +HU’ with ‘siControl +HU’), indicating a greater chance of replication fork slowing, pause, or collapse during the second (IdU) labelling period in cells depleted for SAF-A. (Note that forks pausing or collapsing in the CldU labelling period will not be counted, since as they produce only CldU labelling they cannot be distinguished from termination sites). This result suggests that SAF-A is required to support processive DNA synthesis under replication stress.

The molecular clamp PCNA is required for processive DNA synthesis by DNA Polymerase δ, and potentially by DNA Polymerase ε as well (Eissenberg et al., 1997). To test whether reduced DNA synthesis processivity in cells depleted for SAF-A (Fig 5B) reflects altered PCNA function in DNA replication, we examined the abundance of chromatin-associated PCNA (Fig 5C). The abundance of the chromatin-associated PCNA appears noticeably reduced in cells depleted for SAF-A compared with the control cells (compare lane 3 & 4). This apparent reduction could reflect altered PCNA loading or unloading on chromatin, or potentially increased modification that alters PCNA gel mobility. We did notice that the proportion of apparently SUMO-modified PCNA within the chromatin-associated fraction was increased in siSAF-A (Fig 5C lane 4, upper PCNA band), compared to Control (Fig 5C lane 4) (Gali et al., 2012). Depletion of SAF-A therefore appears to impact composition of replication machinery, in a way that may explain the slow fork speed and reduced DNA synthesis processivity in cells depleted for SAF-A.

The nascent DNA labelling experiments demonstrate that SAF-A is required for robust replication fork progression, as well as to support origin licensing.

### SAF-A affects the replication timing of domain boundaries

Given its effect on replication origin activation and fork progression, we examined whether SAF-A mediates normal DNA replication timing. To allow the detection of any changes that might not be evident in a population analysis (e.g. increased variability in time of replication that does not affect the average value), we examined the replication timing programme in single cells using a recently described method (Miura et al., 2020; Takahashi et al., 2019). Briefly, in this method single mid-S phase cells are collected by cell sorting based on their DNA content, then NGS library preparation and copy number sequencing carried out for each individual single cell. As a control, we carried out similar analysis using a pool of 100 mid-S cells. The relative copy number of 200 kb segments was calculated based on the number of NGS reads, normalised against reads obtained from G1 cells.

We compared the replication timing profiles of 33 single mid-S hTERT-RPE1 cells for siControl, and 25 single mid-S cells for siSAF-A (Fig 6A). While overall replication timing profiles are largely conserved in siSAF-A cells, we found that the boundaries of the replication timing domains are less uniform in siSAF-A cells. For example, in the regions shown magnified at the bottom of Fig 6A, the siControl cells show clear boundaries between unreplicated (blue) and replicated (red) domains, with a fairly uniform pattern across the 33 analysed cells. In siSAF-A cells in contrast, the boundary position shows more variation between single cells, resulting in a lack of clear boundaries when viewed across the population. Statistical comparison of single-cell replication timing between siControl and siSAF-A cells confirms this notion (see “−log_10_P plot in Fig 6A). This finding is consistent with the proposed function of SAF-A in defining chromatin domain boundaries (Fan et al., 2018).

**Figure 6.**
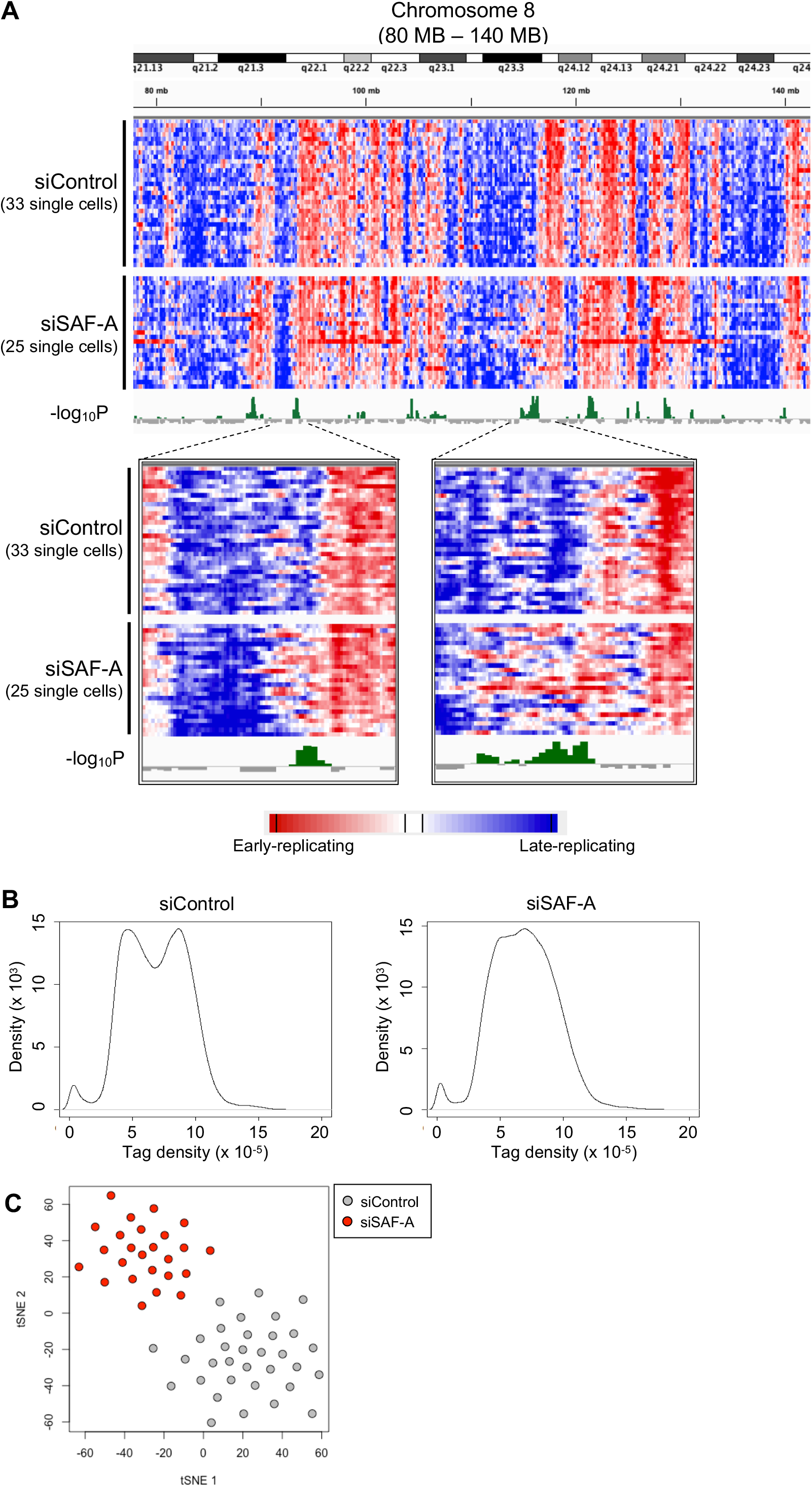
Replication timing is affected by SAF-A. (A) 60 Mbase region of Chromosome 8 illustrating the impact of depleting SAF-A on single-cell replication timing profiles. Heat maps show replication in single mid-S phase cells (red: early replicating, blue: late replicating). Each horizontal line represents the replication timing of a single cell (33 siControl cells and 25 siSAF-A cells), in 200-kb windows. The “-log_10_P” plot (green) shows statistical significance of the difference between single-cell replication timing of siControl and siSAF-A cells. Two specimen regions showing differences between siControl and siSAF-A are shown magnified at the bottom. Note that horizontal streaks in blue or red indicate gain or loss of chromosomal segments, so do not represent timing. (B) Distribution of NGS tag density in 100 cells of siControl and siSAF-A. One hundred mid-S cells were collected by a cell sorter, and NGS libraries were prepared. Tag densities were calculated for 200 kb sliding windows at 40 kb intervals across the genome. (C) t-SNE clustering analysis of replication timing in siControl and siSAF-A cells.

To quantitatively confirm the variation in the position of domain boundary in individual siSAF-A cells, we compared the collective distribution of NGS reads per 200 kb sliding window (=tag density) at 40-kb intervals (Miura et al., 2020). In the pool of 100 mid-S cells from the siControl, distribution of the tag density forms two overlapped peaks (Fig 6B left), representing unreplicated (left peak) and replicated (right peak) portions of the genome. The separation of these peaks in the 100 pooled cells means that (1) unreplicated and replicated domains are distinct in each cell and (2) this distinct pattern is essentially conserved in the 100 cells. In other words, the replication timing programme is well conserved in these 100 cells in siControl cells. In contrast, tag density from the 100 mid-S siSAF-A cells does not show clear peak separation (Fig 6B right). Note however that we do see clear separation of two peaks when analysing single siSAF-A cells at mid-S (Fig S4), as for single siControl cells, indicating that unreplicated and replicated domains are effectively distinguished in analysis of single cells. Therefore, the poor peak separation of tag density in the 100 mid-S siSAF-A cell pool is due to poor conservation of the replication timing programme between single cells.

Although the overall replication timing profiles appear fairly similar in the Fig 6A heat maps, t-SNE clustering analysis (van der Maaten, 2014; van der Maaten and Hinton, 2008) of the distribution of early and late domains in single cells showed a clear separation of the siControl and siSAF-A populations (Fig 6C), indicating that genome-wide replication timing program is indeed altered in siSAF-A cells.

Taken these observations together, we conclude that in cells depleted for SAF-A, the genome-wide DNA replication timing programme is less well-defined, becoming more ‘blurred’ and unstable particularly at domain boundaries.

### SAF-A prevents spontaneous quiescence

Our data suggest that DNA replication is aberrant at various stages in cells depleted for SAF-A, even without exogenous replication stress (Fig 1F, 1G, S1E, and 5A). Recent studies suggest that cells with incomplete DNA replication and/or DNA damage can progress through mitosis but may activate the p53-mediated G1 checkpoint in the subsequent cell cycle, leading to a transient quiescence of daughter cells (Arora et al., 2017; Barr et al., 2017). Such delayed progression can be monitored by the expression of a CDK inhibitor p21^WAF1^. We tested the possibility that replication problems in siSAF-A cells leads to spontaneous quiescence, by looking at the expression of p21. Cells depleted for SAF-A show clear expression of p21 without any exogenous damage (Fig 7A), whereas the expression of p21 is barely detectable in siControl cells. We next tested the distribution of p21-positive cells in the cell cycle using flow cytometry (Fig 7B). As expected, a significant proportion of siSAF-A cells with a ‘G1 phase’ DNA content show p21 expression, suggesting these cells are in quiescence (= G0 phase). Interestingly, in cells depleted for SAF-A, a fraction of G2 cells already express p21. Similar expression of p21 in G2 cells is reported for cells with DNA damage before entering quiescence (Arora et al., 2017; Barr et al., 2017).

**Figure 7:**
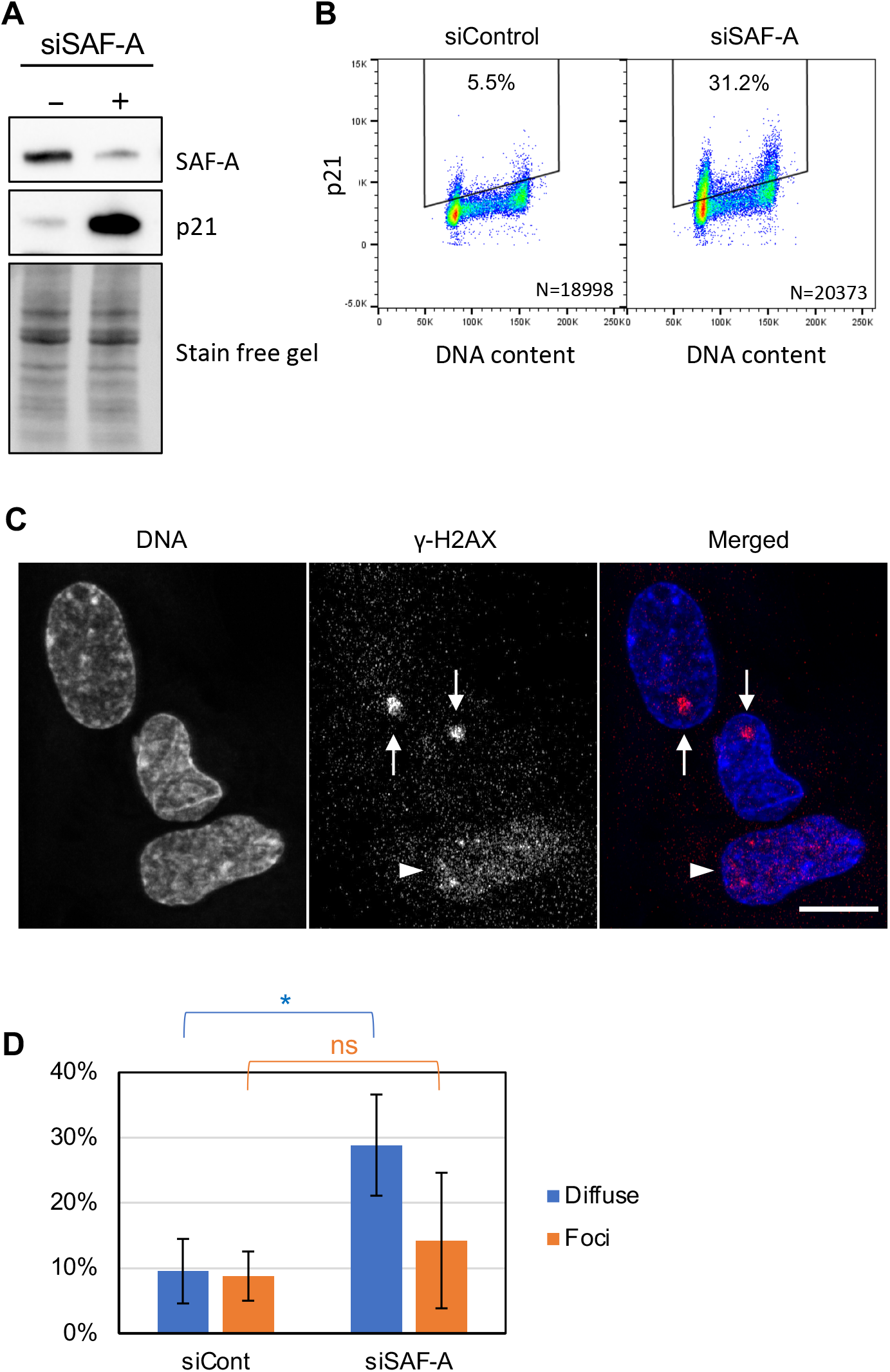
Loss of SAF-A leads to spontaneous replication stress and quiescence. (A) Depletion of SAF-A leads to p21 expression. Whole-cell extracts were prepared from cells treated with control siRNA (−) and SAF-A siRNA (+), and the abundance of SAF-A and p21 was examined by the western blotting. Stain-free gel image is shown as the loading control. (B) Cell cycle analysis of p21 expression. Cells treated with the control siRNA (siControl) and SAF-A siRNA (siSAF-A) were analysed for their DNA content and p21 expression by flow cytometry. Gates used to identify p21-positive cells are shown. (C) Specimen image showing different γ-H2AX localisation patterns. Arrowheads indicate cell with ‘diffuse’ γ-H2AX localisation, and arrows indicate cells with γ-H2AX foci. Scale bar is 10 μm. (D) Depletion of SAF-A leads to spontaneous replication stress. Cells treated with control siRNA (siControl) and SAF-A siRNA (siSAF-A) were analysed for the localisation of γ-H2AX by immunofluorescence. Percentage of cells with either γ-H2AX foci or diffuse γ-H2AX localisations were scored. Averages and standard deviations from 4 independent experiments are shown. At least 50 cells were analysed for each condition. The p-values were calculated by Student’s t-test. ns, not significant; *p ≤ 0.05.

However, it is unclear whether SAF-A depletion itself leads to DNA damage. A previous study demonstrated that depletion of SAF-A increased the proportion of cells showing diffuse localisation of the histone variant H2AX phosphorylated at its C-terminus (called γ-H2AX) (Nozawa et al., 2017). Although γ-H2AX has been commonly used as a DNA damage marker, recent studies suggest that diffuse localisation of γ-H2AX within the nucleus is indicative of replication stress rather than DNA damage, whereas a focal γ-H2AX localisation pattern represents DNA damage (Dhuppar et al., 2020; Moeglin et al., 2019). We assessed the impact of depletion of SAF-A with or without replication stress based on γ-H2AX localisation pattern. Fig 7C shows a specimen image with ‘diffuse’ and ‘focal’ γ-H2AX localisation patterns. Control cells without replication stress show few cells with either γ-H2AX pattern, the majority of cells showing no apparent γ-H2AX signal (Fig 7D, siCont). However, replication stress (induced by 3 hr HU treatment) significantly increased the proportion of cells with ‘diffuse’ γ-H2AX (29%; Fig S5), consistent with the suggestion that diffuse γ-H2AX signals represent DNA replication stress. In contrast, 25% of cells depleted for SAF-A have diffuse γ-H2AX even without HU treatment (Fig 7D, siSAF-A), suggesting that depletion of SAF-A imposes replication stress on cells. In a separate experiment, we confirmed that virtually all cells with diffuse γ-H2AX signal are in S phase (Fig S5B), and that a large fraction of S phase cells have diffuse γ-H2AX signal when SAF-A is depleted (Fig S5C).

These data demonstrate that cells depleted for SAF-A suffer from constant replication stress, leading to more frequent (or more extended) quiescence than in control cells, which can at least partly explain the slower cell proliferation (Fig 1C) and a reduced fraction of S-phase cells (Fig 1F).

Taken all these together, our data demonstrate that SAF-A supports DNA replication by promoting origin licensing, fork progression speed, and fork processivity, probably by modulating the chromatin compaction to ensure the optimal structure for robust DNA replication.

## Discussion

Our investigation of effects of SAF-A on DNA replication establishes that SAF-A promotes replication licensing (Figs 2 & 3). Consistent with this effect, cells depleted for SAF-A showed increased origin spacing when compared to control cells, as well as reduced ability to activate dormant origins under replication stress (Fig 4C). In cells depleted for SAF-A, origin licensing is more sensitive to chromatin compaction induced by sucrose treatment (Fig 2C), suggesting that SAF-A counteracts over-compaction to support origin licensing. Conversely, it was recently demonstrated that the histone methyltransferase SET8 limits replication licensing (Shoaib et al., 2018), presumably through its activity in histone H4K20 methylation. It therefore appears that the correct level of origin licensing requires an appropriate balance between oppositely acting cellular mechanisms that specify chromatin compaction.

Once origins have initiated, replication fork progression is also affected by SAF-A depletion, with fork speed significantly reduced (Fig 5A). This reduced fork speed may potentially reflect an increased incidence of ‘chromatin obstacles’ in the absence of SAF-A, corresponding to hard-to-replicate sites that challenge the replication machinery (Gadaleta and Noguchi, 2017). It has been demonstrated that processive replication through heterochromatin regions is coupled with local chromatin decompaction (Chagin et al., 2019), so that chromatin over-compaction in the absence of SAF-A may increase the number of replication fork impediments, causing the reduced fork processivity observed (Fig 5B). We note also that chromatin-associated PCNA is reduced in cells depleted for SAF-A (Fig 5C), which may be associated with reduced processivity.

Despite having a moderate effect on origin licensing, depletion of SAF-A has a fairly mild effect on EdU incorporation levels without additional replication stress (Fig 1F, 1G, and S1E). This probably reflects the fact that under normal circumstances, MCM complex is loaded at a larger number of sites than will be utilised, so that a modest reduction in origin licensing has only slight impact on cellular DNA replication dynamics under unperturbed conditions (Ge et al., 2007; Woodward et al., 2006). We find however that there is a stronger requirement for SAF-A in enabling cells to recover from replication stress (Fig 1D, 1E, S1C, and S1D). One possible explanation, based on the origin licensing defect of SAF-A-depleted cells, is that insufficient ‘dormant’ origins are available for activation to enable proper recovery from stress (Fig 4C). Inadequate licensing that fails to provide enough dormant origins may lead to chromosome segments remaining unreplicated, if incoming replication forks from both directions collapse under replication stress conditions (McIntosh and Blow, 2012).

As discussed above, it is also possible that SAF-A affects the stability of replication forks through its effect on chromatin compaction. In the absence of SAF-A replication forks may collapse into a conformation that is harder to restart, especially given the low levels of PCNA on chromatin. Intriguingly, SAF-A is also proposed to be involved in double-strand-break damage repair (Hegde et al., 2016)., hinting that there might be a common role for SAF-A in modulating chromatin at stalled replication forks and break repair sites.

Although depletion of SAF-A leads to reduced overall licensing and increased origin spacing, we find that the genome-wide replication timing programme is not severely affected (Fig 6A). This is perhaps consistent with the fact that each replication timing domain contains multiple replication origins whose activation is concomitantly regulated. Therefore, even if some licenced origins are lost from a replication timing domain, the domain is likely still to retain its original replication timing program, enabled by correctly regulated initiation at the remaining origins. We did however observe that the sharp boundaries that normally delineate replication timing domains tend to be obscured, with considerably increased cell-to-cell heterogeneity of these boundaries (Fig 6A). Such increased heterogeneity in timing domain boundaries will contribute to the poor separation of ‘early’ and ‘late’ replication peaks in the analysis of genome-wide tag density distribution (Fig 6B), and is likely to be an important parameter in the separation of siControl cells and siSAF-A cells by t-SNE analysis (Fig 6C). Interestingly, SAF-A protein has been reported to interact with chromatin domain boundary proteins including CTCF and cohesion subunit RAD21 (Fan et al., 2018; Zhang et al., 2019) and has been reported to in involved in defining chromatin domain boundaries (Fan et al., 2018).

SAF-A has been implicated in inactivation of X chromosomes (Lu et al., 2020; Smeets et al., 2014). The hTERT-RPE1 cells used for our timing analysis are female, but we did not observe any obvious impact of depleting SAF-A on X chromosome replication timing (not shown). However, subtle changes in the inactive X chromosome replication timing might not be detected, given our methodology did not separately analyse the two X chromosomes.

RNA appears to be a functional component of chromatin (Brockdorff, 2019; Michieletto and Gilbert, 2019; Rodriguez-Campos and Azorin, 2007), but its molecular contribution is not well understood. One recent study proposes that chromatin-associated RNA promotes open chromatin structures by neutralising the positive charges on histone tails (Dueva et al., 2019). SAF-A could assist this process by tethering RNA molecules in the proximity of chromatin. Our observation of abnormal DNA distribution in SAF-A-depleted nuclei (Fig 1A, 1B and S1A) may be connected to this function. We found that chromatin tends to form abnormal dense clusters in the absence of SAF-A. RNA and splicing machinery are likely to be enriched in the nuclear regions devoid of chromosomal DNA (Smeets et al., 2014). The ‘polarised’ chromatin distribution we observed may reflect a role for SAF-A in the recruitment of chromatin-associated RNA.

A recent high-throughput proteomics study additionally demonstrates that many proteins implicated in DNA replication form large protein complexes depending on RNA association (Caudron-Herger et al., 2019). SAF-A could be envisaged to assist the assembly or activity of such RNA-dependent complexes.

In summary, we have demonstrated that SAF-A is required for robust DNA replication, both in unperturbed conditions and in the recovery from replication stress. Moreover, we show that depletion of SAF-A leads to spontaneous replication stress and increased quiescence. Our observations indicate that SAF-A function is needed for cells to deal with normally arising levels of replication stress and maintain the cell’s potential to proliferate. Expression of SAF-A is increased in a wide range of cancers (The Cancer Genome Atlas), suggesting that cancer cells rely on SAF-A function, and that tumorigenesis-associated events may require increased SAF-A activity (potentially relating to increased replication origin activity in cancer cells (Macheret and Halazonetis, 2018). Replication stress is an important hallmark of cancer (Gaillard et al., 2015; Macheret and Halazonetis, 2015), and an intriguing possibility is that SAF-A is important for cancer cell survival in the context of such replication stress. Conversely, SAF-A loss-of-function alleles are linked to developmental disorders including microcephaly (Durkin et al., 2020; Leduc et al., 2017; Yates et al., 2017). Overall, our findings reported here show that the promotion of robust DNA replication by SAF-A is crucial for its role in supporting cellular capacity for proliferation.

## Materials and methods

### Cell lines

Cell lines hTERT-RPE1 (Bodnar et al., 1998) and HEK293 FLAG-ORC1 were as previously described (Tatsumi et al., 2003).

### Cell culture

All human cell lines were cultivated in synthetic defined media (described below) supplemented with 10% foetal bovine serum (tetracycline-free), 100 U/ml penicillin, and 100 μg/ml streptomycin in 5% CO_2_, ambient O_2_ and at 37°C. hTERT-RPE1 cells were cultivated in DMEM (Gibco) or DMEM-F12 (Gibco) where specified. Other cell lines were cultivated in DMEM.

### siRNA

siRNA used are:

SAF-A siRNA - Human HNRNPU (3192) ON-TARGETplus SMARTpool from Dharmacon Cat#L-013501-00-0005

Control siRNA - Luciferase (GL2) from Dharmacon Cat#D-001100–01

### Antibodies

Primary antibodies used were;

SAF-A - Mouse monoclonal [3G6] (Abcam, ab10297)

FLAG - FLAG tag antibody [M2] (Sigma, F-1804)

MCM3 - MCM3 Antibody (N-19) goat polyclonal IgG (Santa Cruz, sc-9850)

CDT1 – Anti-CDT1/DUP antibody [EPR17891] rabbit monoclonal (Abcam, ab202067)

CDC6 - Rabbit monoclonal [EPR714(2)] (Abcam, ab109315)

ORC1 – ORC1 Antibody [F-10] mouse monoclonal IgG1 (Santa Cruz, sc398734) ORC2 – rabbit polyclonal (Bethyl, A302-734A)

p21 – p21 Antibody (C-19) rabbit polyclonal (Santa Cruz, sc397)

PCNA – mouse monoclonal IgG2a (Santa Cruz, sc-56)

H3 – Rabbit polyclonal (Abcam, ab1791)

CldU - Rat monoclonal anti-BrdU [BU1/75 (ICR1)], (Abcam, ab6326)

IdU - Mouse monoclonal anti-BrdU (BD Biosciences, Cat# 347580)

ssDNA - mouse monoclonal IgG3 (Millipore MAB3868)

γ-H2AX antibodies used were p-Histone H2A.X S139 (20E3) Rabbit mAb (Cell Signalling Technology, #9718) and Alexa Fluor 647 Mouse anti-H2AX (pS139) Clone N1-431 (BD Pharmingen, 560447).

Secondary antibodies used were:

AlexaFluor647 Donkey anti-rabbit IgG (H+L) (Abcam, ab150063)

AlexaFluor647 anti-goat IgG (H+L) (Abcam, ab150135)

AlexaFluor488 anti-mouse IgG (H+L) (Abcam, ab150177)

Alexa Fluor 594 anti-rat IgG (H+L) (Molecular Probes A-11007)

Alexa Fluor 350 Goat anti-Mouse IgG (H+L) (Molecular Probes A-11045)

Alexa Fluor 488 anti-mouse IgG1 (Molecular Probes A-21121)

### Chromatin fractionation

To prepare chromatin-enriched fractions for analysis of PCNA and ORC2 (Fig 5C and S2C, respectively), cells were lysed in cytoskeleton (CSK) buffer (10 mM HEPES-KOH [pH 7.4], 100 mM NaCl, 3 mM MgCl2, 300 mM sucrose), containing 0.2% Triton X-100, 1X EDTA-free protease inhibitor (Roche, 04693159001) and 1X HALT protease and phosphatase inhibitor (Thermo Scientific, 78446) for 10 min on ice. Lysed cells were then centrifuged for 3 min at 2000xg. The pellet was washed once with CSK buffer, centrifuged for 4 min at 3200 rpm, and resuspended in CSK buffer containing 10 μl/ml Benzonase for 30 min on ice. Samples were boiled in 1X Laemmli sample buffer for 10 min and 5% β-mercaptoethanol was added.

To prepare chromatin enriched fractions for analysis of p21 (Fig 7a), CDT1 (Fig S2C left), CDC6 and ORC1 (Fig S2C middle), cells were lysed in Low Salt Extraction (LSE) buffer (10mM K-phosphate [pH 7.4], 10 mM NaCl, 5 mM MgCl2) containing 0.1% Igepal CA-630 and 1mM PMSF for 5 min on ice. Lysed cells were then centrifuged and the pellet was washed once with LSE buffer. The pellet was resuspended and boiled in 1X Laemmli sample buffer for 10 min and 5% β-mercaptoethanol was added.

Protein concentrations in whole cell extracts were determined using the Bio-Rad RC DC Protein assay kit. For Western blots, an equal amount of total protein was loaded on each whole cell extract lane, and the loading for the chromatin fractions was then determined based on cell-equivalency. Equal loading was further confirmed by examining total protein using Bio-Rad stain-free gels.

### DNA combing

For analysis of nascent DNA on DNA fibres, cells were pulse-labelled sequentially with CldU and IdU for 20 min each. Cells were then collected and DNA combing carried using FiberComb instrument (Genomic Vision) according to the manufacturer’s instructions. Detection of CldU and IdU was as previously described (Garzon et al., 2019). Images were acquired on Zeiss Axio Imager M2 microscope and 63x/NA1.4 objective equipped with ORCA-Flash 4.0LT CMOS camera (Hamamatsu Photonics). Images were analysed as previously described (Garzon et al., 2019). For inter-origin distance measurements, 1 μm was converted to 2 kb based on a predetermined value (Bensimon et al., 1994; Bensimon et al., 1995).

### Flow cytometry

Cell cycle analysis of cells stained with DAPI was performed as described (Hiraga et al., 2017; Watts et al., 2020). EdU labelling and its detection by flow cytometry have been previously described (Hiraga et al., 2017). Detection and analysis of chromatin-bound proteins by flow cytometry were performed as previously described (Hiraga et al., 2017) with multiplexing as described below. Data were acquired on Becton Dickinson LSRII or Fortessa flow cytometers with FACSDiva software, and analysed using FlowJo software (Ver. 10.4.2).

We found that apparent MCM levels per cell are very sensitive to the number of cells analysed (i.e. the ratio of cells to antibody during immunostaining), causing tube-to-tube variations. To avoid this issue, we adopted a “multiplexing” strategy. In brief, before immunostaining, samples were differentially labelled with CellTrace Yellow (Molecular Probes) at a concentration unique to each sample (between 0 μM and 0.5 μM final concentration). Differentially stained samples were then mixed, and immunostained in a single tube, to eliminate tube-to-tube variations. After data acquisition by flow cytometry, cell populations were separated based on their CellTrace Yellow signal levels. We confirmed that the CellTrace Yellow signal does not affect the quantification of AlexaFluor 488 and AlexaFluor 647 signals.

### Microscopy

For visualisation of chromatin DNA within the nucleus, cells were grown on ibidi chambered slides (ibi-treated). Cells were washed with PBS, and fixed with neutral buffered 4% formaldehyde (Sigma) for 15 min at RT, then permeabilised with 0.1% Triton X-100 in PBS for 15 min at RT. After washing cells three times with PBS + 0.1% Igepal CA-630, DNA was stained with 0.25 μg/ml DAPI for 30 min. Cells were finally washed and mounted in ibidi mounting medium. Eleven Z-section images were acquired at 170 nm intervals on Zeiss LSM-880/AiryScan microscope with 63x/NA 1.3 objective. The middle section of the Z-stacks was assigned as the plane where each nucleus has the largest XY projection. After AiryScan processing (with the automatic 3-D AiryScan processing condition), the middle section was used for analysis. The areas of DAPI-positive and DAPI-negative regions were determined in an unbiased manner by a custom pipeline utilising Minimal Cross-Entropy on CellProfiler 3.19 (McQuin et al., 2018).

For visualisation of EdU incorporation and immunofluorescence detection of γ-H2AX, cells were grown and fixed as above, and kept in 70% ethanol. Cells were then washed with PBS, permeabilised with 0.5% Triton X-100 in PBS, and incorporated EdU was visualised using Alexa Fluor 488 EdU imaging kit (Molecular Probes C10337) according to the manufacturer’s instruction, followed by indirect immunofluorescence staining of γ-H2AX. Antibodies used were p-Histone H2A.X S139 (20E3) Rabbit mAb (Cell Signalling, #9718) and AlexaFluor 647 anti-rabbit IgG (Abcam, ab150063). Z-stack images were acquired at 250 nm intervals to cover entire nuclei in the field. After AiryScan processing, maximum intensity z-projection images were created for downstream analysis using ImageJ. Detection of cells with diffuse γ-H2AX or γ-H2AX foci were carried out by using a custom CellProfiler pipeline.

### Single-cell replication timing analysis and bioinformatics

Single-cell replication timing of siControl and siSAF-A hTERT-RPE1 cells were analysed as described (Miura et al., 2020; Takahashi et al., 2019). The “−log_10_P” values were calculated by comparing the distribution of single-cell replication timing of 100-kb segments between siControl and siSAF-A cells using student’s t-test. t-SNE clustering analysis of replication timing profile data was performed using Rtsne R library (Krijthe, 2015), with R version 3.6.1. Homo sapiens (human) genome assembly GRCh38 (hg38) from Genome Reference Consortium was used throughout the analysis.

## Acknowledgements

Information for SAF-A expression was obtained at The Cancer Genome Atlas (TCGA) Research Network (https://www.cancer.gov/tcga). We thank Dr Ryu-suke Nozawa for help in the early stage of the project, and Professor Julian Blow for advice on the 3D licensing assay. Thanks to the staff of the Iain Fraser Cytometry Centre and Microscopy and Histology facility at the University of Aberdeen. CC was supported by a BBSRC EASTBIO Doctoral Training programme PhD studentship. SH was supported by Daiwa Anglo-Japanese Foundation (12928/13746). Work in the Hiraga-Donaldson lab supported by Cancer Research UK awards C1445/A19059 and DRCPGM\100013. NG is supported by Medical Research Council (MC_UU_00007/13).

## Supplementary Figure Legends

**Figure S1: SAF-A is required for robust DNA replication**

(A) The ratio of “DAPI-positive” areas per nucleus was measured as in Fig 1B (repeat experiment). Median with 95% confidence interval shown in red. (B) DNA content analysis of asynchronously growing control siControl- and siSAF-A-treated cells analysed by flow cytometry. (C) Impact of SAF-A depletion on recovery from replication inhibition. Cells were arrested by HU and released as in Fig 1D & E, and changes in DNA content and EdU incorporation analysed at times indicated. Gates were set as indicated, to account for shift in basal EdU signal over time. (D) EdU incorporation in EdU-positive populations in control and siSAF-A cells. Cells were treated as described in Fig 1D & E, and the amount of EdU incorporated in each EdU-positive cell was analysed by flow cytometry at indicated time points. Note that this measurement is based on only EdU-positive cells identified as in (C), and is unaffected by the EdU-negative populations. Violin plots show the median (solid line) and quartiles (dotted lines). As distribution was not normal, the p-value was calculated by Mann-Whitney-Wilcoxon test. (E) EdU incorporation in cells at various stages within S phase. The S phase cells were gated into early-, mid-, and late-S phase populations based on DNA content (shown in D), and EdU incorporation measured in each population. The p-value was calculated by Mann-Whitney-Wilcoxon test. ns, not significant, ***p ≤ 0.001, ****p ≤ 0.0001.

**Figure S2: Impact of SAF-A depletion on replication licensing**

(A) HEK293 FLAG-ORC1 cells were pretreated with sucrose before collection as in Fig 2C. The abundance of chromatin-associated CDT1 protein was analysed by multiplexed flow cytometry. Note that these samples were 4-way multiplexed and analysed together, so that chromatin association of CDT1 can be accurately compared across all four panels. Contour lines set to 5%. (B) SAF-A depletion does not affect chromatin association of ORC2. Chromatin association of ORC2 was assessed by flow cytometry in hTERT-RPE1 cells as in Fig 3A. Contour lines set to 5%. (C) Impact of SAF-A depletion on replication licensing was confirmed by western analysis of chromatin-enriched fractions of hTERT-RPE1 cells. Stain-free gel was used as a loading control.

**Figure S3: DNA synthesis during mild HU treatment**

Cells were treated as described in Fig 4B, and IdU tract lengths were measured as a proxy of DNA replication fork speed. As distribution not normal, statistical significance was tested by Mann-Whitney-Wilcoxon test.

ns, not significant; *p ≤ 0.05.

**Figure S4: Tag distribution in single cells**

Tag density distributions in 4 single siControl and 4 single siSAF-A cells. Tag densities were calculated for 200 kb sliding windows at 40 kb intervals over the genome for each cell, as in Fig 6A.

**Figure S5: Diffuse γ-H2AX signal is associated with replication stress**

(A) Replication stress induces diffuse γ-H2AX signal. γ-H2AX was analysed in siControl cells with or without DNA replication stress (3 hr treatment with 1 mM HU), and cells with γ-H2AX signal were scored as in Fig 7D. The p-values were calculated by Student’s t-test. (B) γ-H2AX signals are detected almost exclusively in S phase cells. Cells were pulse-labelled with EdU to detect S phase cells. γ-H2AX signals were detected as in Fig 7D, and the percentage of S phase (= EdU-positive) cells was calculated amongst cells with diffuse γ-H2AX signal. Averages and ranges from two independent experiments are shown. (C) The majority of SAF-A-depleted S phase cells are under replication stress. Percentage of cells with diffuse γ-H2AX signal was calculated within the S phase (=EdU-positive) population. Averages and ranges from two independent experiments are shown.

## References

Arora, M., Moser, J., Phadke, H., Basha, A.A., and Spencer, S.L. (2017). Endogenous Replication Stress in Mother Cells Leads to Quiescence of Daughter Cells. Cell Rep 19, 1351–1364.

Barr, A.R., Cooper, S., Heldt, F.S., Butera, F., Stoy, H., Mansfeld, J., Novak, B., and Bakal, C. (2017). DNA damage during S-phase mediates the proliferation-quiescence decision in the subsequent G1 via p21 expression. Nat Commun 8, 14728.

Bensimon, A., Simon, A., Chiffaudel, A., Croquette, V., Heslot, F., and Bensimon, D. (1994). Alignment and sensitive detection of DNA by a moving interface. Science 265, 2096–2098.

Bensimon, D., Simon, A.J., Croquette, V.V., and Bensimon, A. (1995). Stretching DNA with a receding meniscus: Experiments and models. Phys Rev Lett 74, 4754–4757.

Bianco, J.N., Poli, J., Saksouk, J., Bacal, J., Silva, M.J., Yoshida, K., Lin, Y.L., Tourriere, H., Lengronne, A., and Pasero, P. (2012). Analysis of DNA replication profiles in budding yeast and mammalian cells using DNA combing. Methods 57, 149–157.

Blow, J.J., and Ge, X.Q. (2009). A model for DNA replication showing how dormant origins safeguard against replication fork failure. EMBO Rep 10, 406–412.

Blow, J.J., and Tanaka, T.U. (2005). The chromosome cycle: coordinating replication and segregation. Second in the cycles review series. EMBO Rep 6, 1028–1034.

Bodnar, A.G., Ouellette, M., Frolkis, M., Holt, S.E., Chiu, C.P., Morin, G.B., Harley, C.B., Shay, J.W., Lichtsteiner, S., and Wright, W.E. (1998). Extension of life-span by introduction of telomerase into normal human cells. Science 279, 349–352.

Boteva, L., Nozawa, R.S., Naughton, C., Samejima, K., Earnshaw, W.C., and Gilbert, N. (2020). Common Fragile Sites Are Characterized by Faulty Condensin Loading after Replication Stress. Cell Rep 32, 108177.

Brockdorff, N. (2019). Localized accumulation of Xist RNA in X chromosome inactivation. Open Biol 9, 190213.

Caudron-Herger, M., Rusin, S.F., Adamo, M.E., Seiler, J., Schmid, V.K., Barreau, E., Kettenbach, A.N., and Diederichs, S. (2019). R-DeeP: Proteome-wide and Quantitative Identification of RNA-Dependent Proteins by Density Gradient Ultracentrifugation. Mol Cell 75, 184–199 e110.

Chagin, V.O., Reinhart, B., Becker, A., Mortusewicz, O., Jost, K.L., Rapp, A., Leonhardt, H., and Cardoso, M.C. (2019). Processive DNA synthesis is associated with localized decompaction of constitutive heterochromatin at the sites of DNA replication and repair. Nucleus 10, 231–253.

Cortez, D. (2015). Preventing replication fork collapse to maintain genome integrity. DNA Repair (Amst) 32, 149–157.

Dhuppar, S., Roy, S., and Mazumder, A. (2020). gammaH2AX in the S Phase after UV Irradiation Corresponds to DNA Replication and Does Not Report on the Extent of DNA Damage. Mol Cell Biol 40.

Dimitrova, D.S., and Gilbert, D.M. (1999). The spatial position and replication timing of chromosomal domains are both established in early G1 phase. Mol Cell 4, 983–993.

Dimitrova, D.S., Prokhorova, T.A., Blow, J.J., Todorov, I.T., and Gilbert, D.M. (2002). Mammalian nuclei become licensed for DNA replication during late telophase. J Cell Sci 115, 51–59.

Douglas, P., Ye, R., Morrice, N., Britton, S., Trinkle-Mulcahy, L., and Lees-Miller, S.P. (2015). Phosphorylation of SAF-A/hnRNP-U Serine 59 by Polo-Like Kinase 1 Is Required for Mitosis. Mol Cell Biol 35, 2699–2713.

Dueva, R., Akopyan, K., Pederiva, C., Trevisan, D., Dhanjal, S., Lindqvist, A., and Farnebo, M. (2019). Neutralization of the Positive Charges on Histone Tails by RNA Promotes an Open Chromatin Structure. Cell Chem Biol 26, 1436–1449 e1435.

Durkin, A., Albaba, S., Fry, A.E., Morton, J.E., Douglas, A., Beleza, A., Williams, D., Volker-Touw, C.M.L., Lynch, S.A., Canham, N., et al. (2020). Clinical findings of 21 previously unreported probands with HNRNPU-related syndrome and comprehensive literature review. Am J Med Genet A 182, 1637–1654.

Eissenberg, J.C., Ayyagari, R., Gomes, X.V., and Burgers, P.M. (1997). Mutations in yeast proliferating cell nuclear antigen define distinct sites for interaction with DNA polymerase delta and DNA polymerase epsilon. Mol Cell Biol 17, 6367–6378.

Fackelmayer, F.O., Dahm, K., Renz, A., Ramsperger, U., and Richter, A. (1994). Nucleic-acid-binding properties of hnRNP-U/SAF-A, a nuclear-matrix protein which binds DNA and RNA in vivo and in vitro. Eur J Biochem 221, 749–757.

Fan, H., Lv, P., Huo, X., Wu, J., Wang, Q., Cheng, L., Liu, Y., Tang, Q.Q., Zhang, L., Zhang, F., et al. (2018). The nuclear matrix protein HNRNPU maintains 3D genome architecture globally in mouse hepatocytes. Genome Res 28, 192–202.

Feng, D., Tu, Z., Wu, W., and Liang, C. (2003). Inhibiting the expression of DNA replication-initiation proteins induces apoptosis in human cancer cells. Cancer Res 63, 7356–7364.

Fragkos, M., Ganier, O., Coulombe, P., and Mechali, M. (2015). DNA replication origin activation in space and time. Nat Rev Mol Cell Biol 16, 360–374.

Frigola, J., He, J., Kinkelin, K., Pye, V.E., Renault, L., Douglas, M.E., Remus, D., Cherepanov, P., Costa, A., and Diffley, J.F.X. (2017). Cdt1 stabilizes an open MCM ring for helicase loading. Nat Commun 8, 15720.

Fu, H., Baris, A., and Aladjem, M.I. (2018). Replication timing and nuclear structure. Curr Opin Cell Biol 52, 43–50.

Gadaleta, M.C., and Noguchi, E. (2017). Regulation of DNA Replication through Natural Impediments in the Eukaryotic Genome. Genes (Basel) 8.

Gaillard, H., Garcia-Muse, T., and Aguilera, A. (2015). Replication stress and cancer. Nat Rev Cancer 15, 276–289.

Gali, H., Juhasz, S., Morocz, M., Hajdu, I., Fatyol, K., Szukacsov, V., Burkovics, P., and Haracska, L. (2012). Role of SUMO modification of human PCNA at stalled replication fork. Nucleic Acids Res 40, 6049–6059.

Garzon, J., Ursich, S., Lopes, M., Hiraga, S.I., and Donaldson, A.D. (2019). Human RIF1-Protein Phosphatase 1 Prevents Degradation and Breakage of Nascent DNA on Replication Stalling. Cell Rep 27, 2558–2566 e2554.

Ge, X.Q., Jackson, D.A., and Blow, J.J. (2007). Dormant origins licensed by excess Mcm2-7 are required for human cells to survive replicative stress. Genes Dev 21, 3331–3341.

Gilbert, D.M. (2010). Cell fate transitions and the replication timing decision point. J Cell Biol 191, 899–903.

Gilbert, D.M., Takebayashi, S.I., Ryba, T., Lu, J., Pope, B.D., Wilson, K.A., and Hiratani, I. (2010). Space and time in the nucleus: developmental control of replication timing and chromosome architecture. Cold Spring Harb Symp Quant Biol 75, 143–153.

Hegde, M.L., Dutta, A., Yang, C., Mantha, A.K., Hegde, P.M., Pandey, A., Sengupta, S., Yu, Y., Calsou, P., Chen, D., et al. (2016). Scaffold attachment factor A (SAF-A) and Ku temporally regulate repair of radiation-induced clustered genome lesions. Oncotarget 7, 54430–54444.

Hiraga, S.I., Ly, T., Garzon, J., Horejsi, Z., Ohkubo, Y.N., Endo, A., Obuse, C., Boulton, S.J., Lamond, A.I., and Donaldson, A.D. (2017). Human RIF1 and protein phosphatase 1 stimulate DNA replication origin licensing but suppress origin activation. EMBO Rep 18, 403–419.

Hiratani, I., Ryba, T., Itoh, M., Yokochi, T., Schwaiger, M., Chang, C.W., Lyou, Y., Townes, T.M., Schubeler, D., and Gilbert, D.M. (2008). Global reorganization of replication domains during embryonic stem cell differentiation. PLoS Biol 6, e245.

Kawabata, T., Luebben, S.W., Yamaguchi, S., Ilves, I., Matise, I., Buske, T., Botchan, M.R., and Shima, N. (2011). Stalled fork rescue via dormant replication origins in unchallenged S phase promotes proper chromosome segregation and tumor suppression. Mol Cell 41, 543–553.

Kiledjian, M., and Dreyfuss, G. (1992). Primary structure and binding activity of the hnRNP U protein: binding RNA through RGG box. The EMBO Journal 11, 2655–2664.

Krijthe, J.H. (2015). Rtsne: T-Distributed Stochastic Neighbor Embedding using a Barnes-Hut Implementation.

Lau, E., Chiang, G.G., Abraham, R.T., and Jiang, W. (2009). Divergent S phase checkpoint activation arising from prereplicative complex deficiency controls cell survival. Mol Biol Cell 20, 3953–3964.

Leduc, M.S., Chao, H.T., Qu, C., Walkiewicz, M., Xiao, R., Magoulas, P., Pan, S., Beuten, J., He, W., Bernstein, J.A., et al. (2017). Clinical and molecular characterization of de novo loss of function variants in HNRNPU. Am J Med Genet A 173, 2680–2689.

Lu, Z., Guo, J.K., Wei, Y., Dou, D.R., Zarnegar, B., Ma, Q., Li, R., Zhao, Y., Liu, F., Choudhry, H., et al. (2020). Structural modularity of the XIST ribonucleoprotein complex. Nat Commun 11, 6163.

Macheret, M., and Halazonetis, T.D. (2015). DNA replication stress as a hallmark of cancer. Annu Rev Pathol 10, 425–448.

Macheret, M., and Halazonetis, T.D. (2018). Intragenic origins due to short G1 phases underlie oncogene-induced DNA replication stress. Nature 555, 112–116.

McIntosh, D., and Blow, J.J. (2012). Dormant origins, the licensing checkpoint, and the response to replicative stresses. Cold Spring Harb Perspect Biol 4.

McQuin, C., Goodman, A., Chernyshev, V., Kamentsky, L., Cimini, B.A., Karhohs, K.W., Doan, M., Ding, L., Rafelski, S.M., Thirstrup, D., et al. (2018). CellProfiler 3.0: Next-generation image processing for biology. PLoS Biol 16, e2005970.

Mendez, J., and Stillman, B. (2000). Chromatin association of human origin recognition complex, cdc6, and minichromosome maintenance proteins during the cell cycle: assembly of prereplication complexes in late mitosis. Mol Cell Biol 20, 8602–8612.

Mendez, J., Zou-Yang, X.H., Kim, S.Y., Hidaka, M., Tansey, W.P., and Stillman, B. (2002). Human origin recognition complex large subunit is degraded by ubiquitin-mediated proteolysis after initiation of DNA replication. Mol Cell 9, 481–491.

Michieletto, D., and Gilbert, N. (2019). Role of nuclear RNA in regulating chromatin structure and transcription. Curr Opin Cell Biol 58, 120–125.

Miura, H., Takahashi, S., Shibata, T., Nagao, K., Obuse, C., Okumura, K., Ogata, M., Hiratani, I., and Takebayashi, S.I. (2020). Mapping replication timing domains genome wide in single mammalian cells with single-cell DNA replication sequencing. Nat Protoc 15, 4058–4100.

Moeglin, E., Desplancq, D., Conic, S., Oulad-Abdelghani, M., Stoessel, A., Chiper, M., Vigneron, M., Didier, P., Tora, L., and Weiss, E. (2019). Uniform Widespread Nuclear Phosphorylation of Histone H2AX Is an Indicator of Lethal DNA Replication Stress. Cancers (Basel) 11.

Moreno, A., Carrington, J.T., Albergante, L., Al Mamun, M., Haagensen, E.J., Komseli, E.S., Gorgoulis, V.G., Newman, T.J., and Blow, J.J. (2016). Unreplicated DNA remaining from unperturbed S phases passes through mitosis for resolution in daughter cells. Proc Natl Acad Sci U S A 113, E5757–5764.

Nevis, K.R., Cordeiro-Stone, M., and Cook, J.G. (2009). Origin licensing and p53 status regulate Cdk2 activity during G(1). Cell Cycle 8, 1952–1963.

Nozawa, R.S., Boteva, L., Soares, D.C., Naughton, C., Dun, A.R., Buckle, A., Ramsahoye, B., Bruton, P.C., Saleeb, R.S., Arnedo, M., et al. (2017). SAF-A Regulates Interphase Chromosome Structure through Oligomerization with Chromatin-Associated RNAs. Cell 169, 1214–1227 e1218.

Ohta, S., Tatsumi, Y., Fujita, M., Tsurimoto, T., and Obuse, C. (2003). The ORC1 cycle in human cells: II. Dynamic changes in the human ORC complex during the cell cycle. J Biol Chem 278, 41535–41540.

Richter, K., Nessling, M., and Lichter, P. (2007). Experimental evidence for the influence of molecular crowding on nuclear architecture. J Cell Sci 120, 1673–1680.

Rodriguez-Campos, A., and Azorin, F. (2007). RNA is an integral component of chromatin that contributes to its structural organization. PLoS One 2, e1182.

Sharp, J.A., Perea-Resa, C., Wang, W., and Blower, M.D. (2020). Cell division requires RNA eviction from condensing chromosomes. J Cell Biol 219.

Shoaib, M., Walter, D., Gillespie, P.J., Izard, F., Fahrenkrog, B., Lleres, D., Lerdrup, M., Johansen, J.V., Hansen, K., Julien, E., et al. (2018). Histone H4K20 methylation mediated chromatin compaction threshold ensures genome integrity by limiting DNA replication licensing. Nat Commun 9, 3704.

Shreeram, S., Sparks, A., Lane, D.P., and Blow, J.J. (2002). Cell type-specific responses of human cells to inhibition of replication licensing. Oncogene 21, 6624–6632.

Smeets, D., Markaki, Y., Schmid, V.J., Kraus, F., Tattermusch, A., Cerase, A., Sterr, M., Fiedler, S., Demmerle, J., Popken, J., et al. (2014). Three-dimensional super-resolution microscopy of the inactive X chromosome territory reveals a collapse of its active nuclear compartment harboring distinct Xist RNA foci. Epigenetics Chromatin 7, 8.

Takahashi, S., Miura, H., Shibata, T., Nagao, K., Okumura, K., Ogata, M., Obuse, C., Takebayashi, S.I., and Hiratani, I. (2019). Genome-wide stability of the DNA replication program in single mammalian cells. Nat Genet 51, 529–540.

Tardat, M., Brustel, J., Kirsh, O., Lefevbre, C., Callanan, M., Sardet, C., and Julien, E. (2010). The histone H4 Lys 20 methyltransferase PR-Set7 regulates replication origins in mammalian cells. Nat Cell Biol 12, 1086–1093.

Tatsumi, Y., Ohta, S., Kimura, H., Tsurimoto, T., and Obuse, C. (2003). The ORC1 cycle in human cells: I. cell cycle-regulated oscillation of human ORC1. J Biol Chem 278, 41528–41534.

van der Maaten, L.J.P. (2014). Accelerating t-SNE using Tree-Based Algorithms. Journal of Machine Learning Research 15, 3221–3245.

van der Maaten, L.J.P., and Hinton, G.E. (2008). Visualizing High-Dimensional Data Using t-SNE. Journal of Machine Learning Research 9, 2579–2605.

Watts, L.P., Natsume, T., Saito, Y., Garzon, J., Dong, Q., Boteva, L., Gilbert, N., Kanemaki, M.T., Hiraga, S.I., and Donaldson, A.D. (2020). The RIF1-long splice variant promotes G1 phase 53BP1 nuclear bodies to protect against replication stress. Elife 9.

Woodward, A.M., Gohler, T., Luciani, M.G., Oehlmann, M., Ge, X., Gartner, A., Jackson, D.A., and Blow, J.J. (2006). Excess Mcm2-7 license dormant origins of replication that can be used under conditions of replicative stress. J Cell Biol 173, 673–683.

Yates, T.M., Vasudevan, P.C., Chandler, K.E., Donnelly, D.E., Stark, Z., Sadedin, S., Willoughby, J., Broad Center for Mendelian, G., study, D.D.D., and Balasubramanian, M. (2017). De novo mutations in HNRNPU result in a neurodevelopmental syndrome. Am J Med Genet A 173, 3003–3012.

Zhai, Y., Cheng, E., Wu, H., Li, N., Yung, P.Y., Gao, N., and Tye, B.K. (2017a). Open-ringed structure of the Cdt1-Mcm2-7 complex as a precursor of the MCM double hexamer. Nat Struct Mol Biol 24, 300–308.

Zhai, Y., Li, N., Jiang, H., Huang, X., Gao, N., and Tye, B.K. (2017b). Unique Roles of the Non-identical MCM Subunits in DNA Replication Licensing. Mol Cell 67, 168–179.

Zhang, L., Song, D., Zhu, B., and Wang, X. (2019). The role of nuclear matrix protein HNRNPU in maintaining the architecture of 3D genome. Semin Cell Dev Biol 90, 161–167.

Zimmerman, K.M., Jones, R.M., Petermann, E., and Jeggo, P.A. (2013). Diminished origin-licensing capacity specifically sensitizes tumor cells to replication stress. Mol Cancer Res 11, 370–380.

