## Supplementary figures and images for "SAF-A promotes origin licensing and replication fork progression to ensure robust DNA replication"

### Supplementary Figures

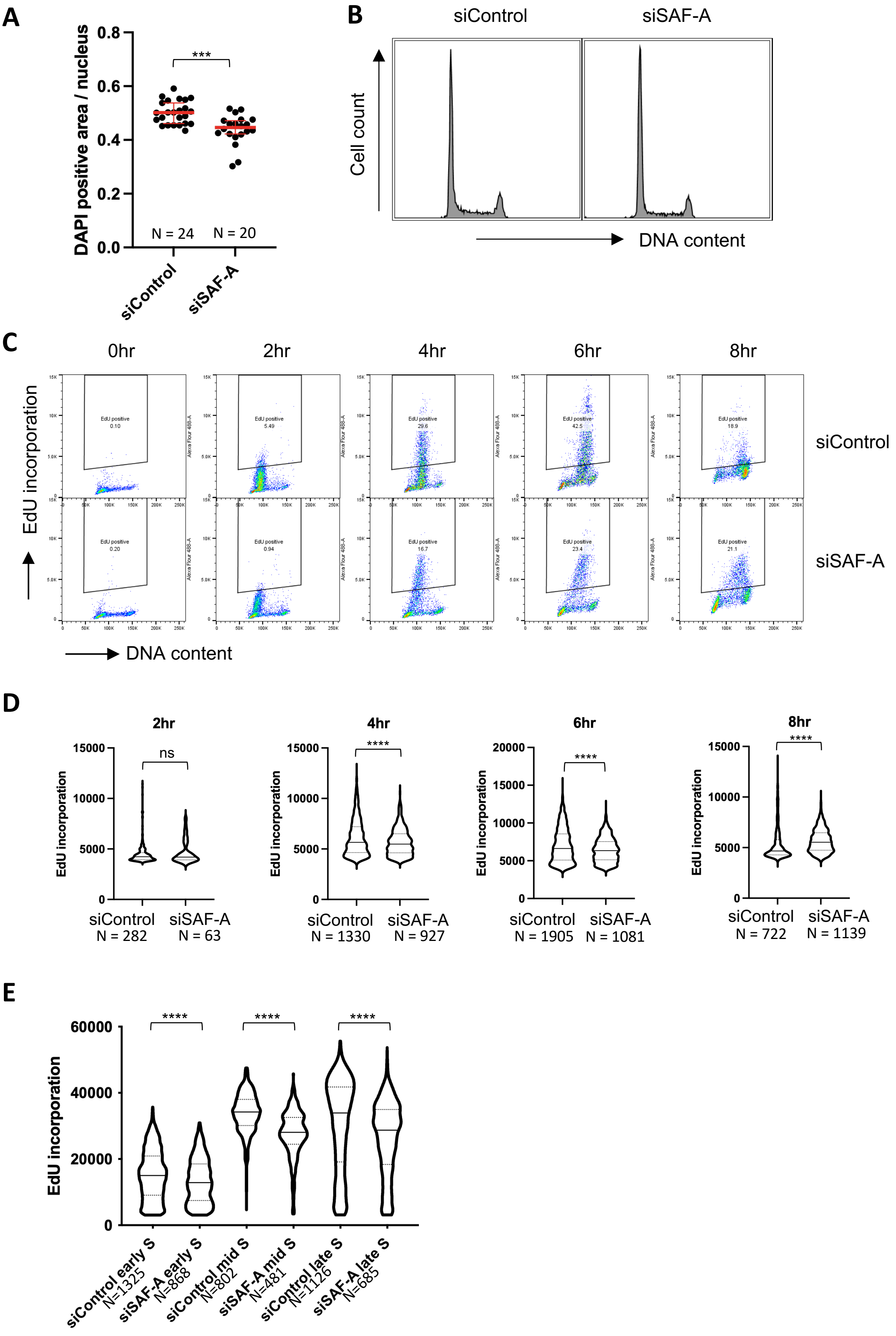

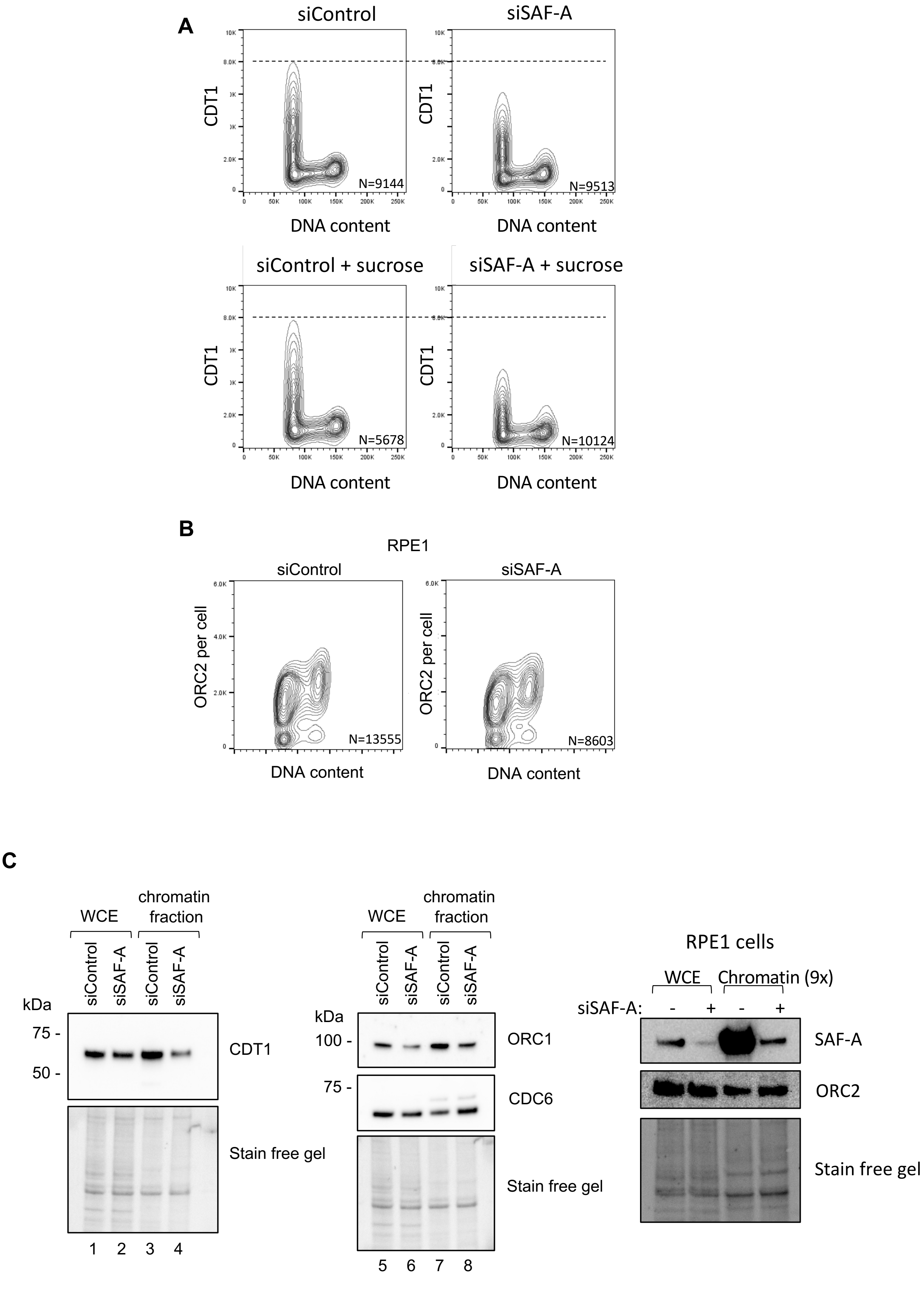

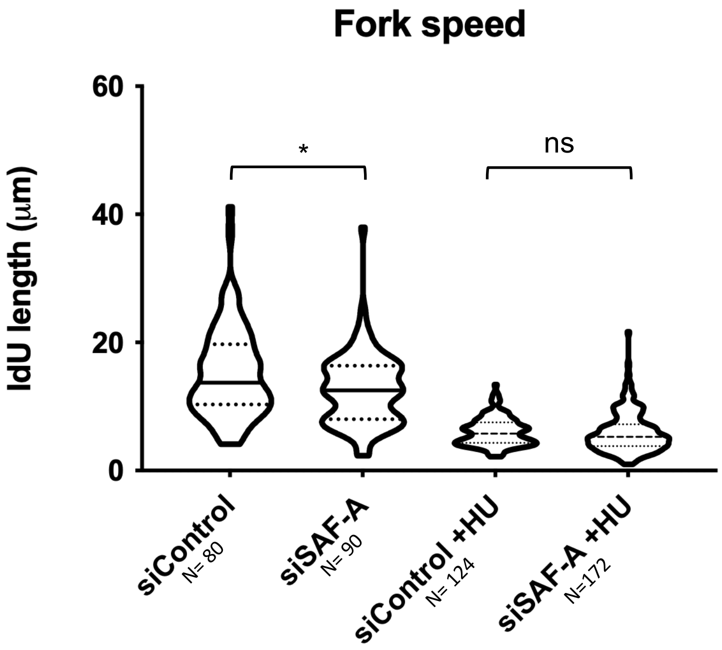

siControl

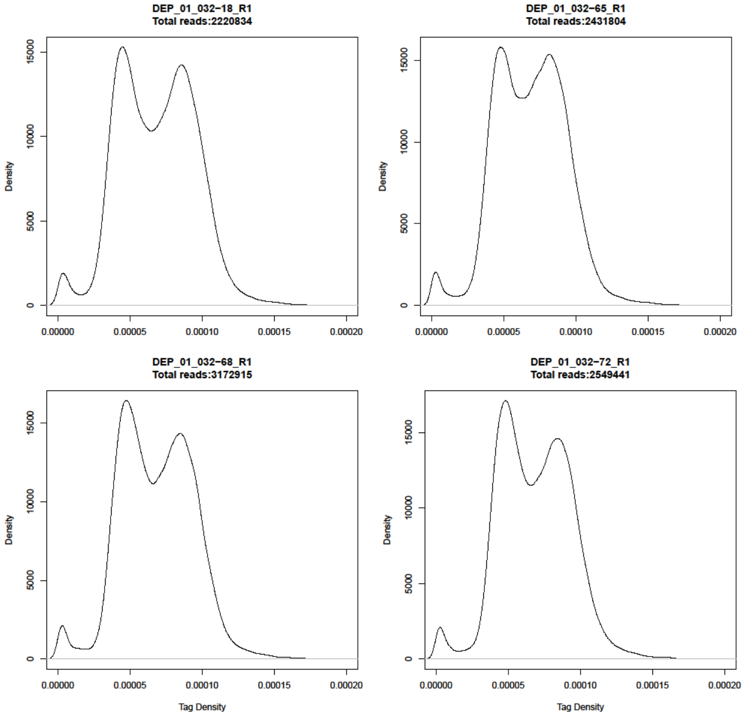


siSAF-A

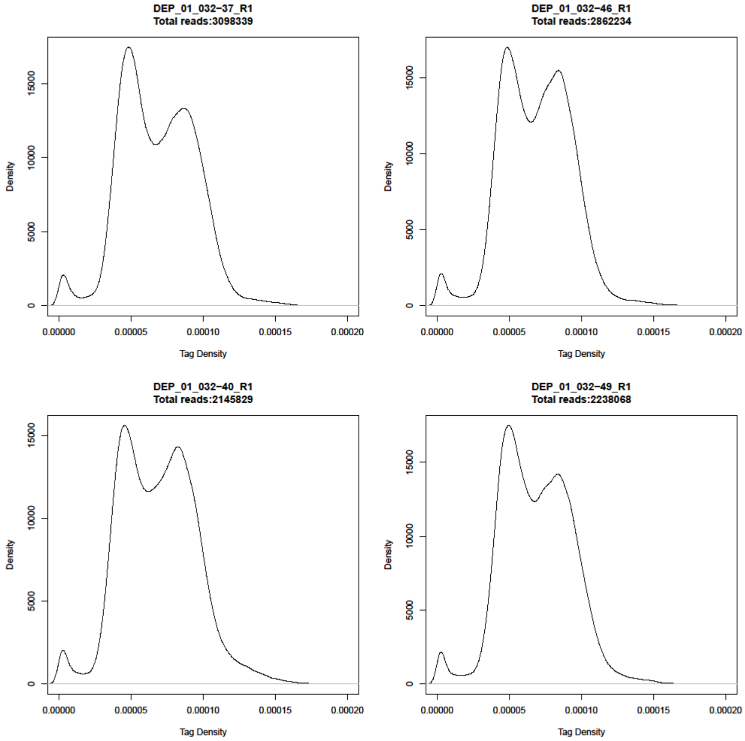


Fig S4

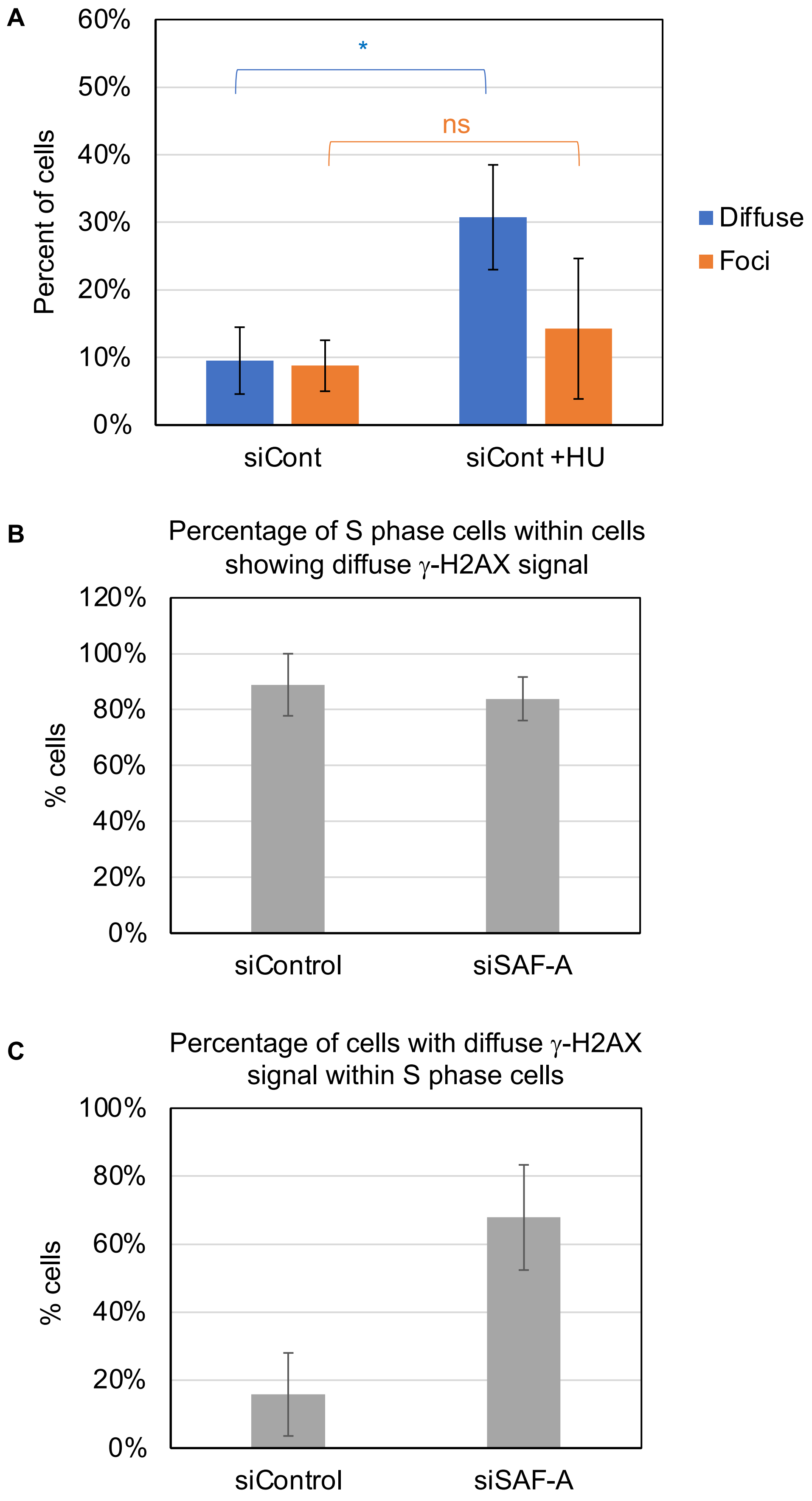
